# Cosmopolitan gut bacteriophages expand the phenotype of health-related bacteria

**DOI:** 10.64898/2026.05.08.723808

**Authors:** Piotr Rozwalak, Andrzej Zielezinski, Maria Alejandra de Angel Fontalvo, Simon Ménard, Sandrine Auger, Bas E. Dutilh

## Abstract

Metagenomics has vastly expanded our knowledge of the human gut virome, yet the focus on dominant taxa has missed low-abundant but highly prevalent bacteriophages. Here, high-sensitivity taxonomic profiling of 53,976 samples from 42 countries, including non-industrialised populations and ancient DNA from five archaeological sites, reveals that only 0.3% of 64,238 phage genera are cosmopolitan, occurring in >20% of all subjects and on six continents. The most prevalent of these is *Mushuvirus*, with two distinct species detected in 60% of individuals, previously missed due to artefacts in reference genomes and their consistently low relative abundance (<0.8%). Population genomics shows that while one species is globally ubiquitous, the other predominates in individuals with West Eurasian ancestry. Together with 12 additional novel genera, we propose to classify these viruses into a new family *Mushuviridae*, found in ∼89% of studied humans. These transposable phages integrate into diverse hosts from two taxonomic orders of short-chain fatty acid–producing bacteria. Multi-omics experiments with *Faecalibacterium* demonstrate that hosts produce and secrete *Mushuvirus*-encoded receptor-binding proteins, which are actively diversified in lysogenic hosts. Together, these findings illuminate a globally distributed, active phage lineage that has persisted for millennia and continues to reshape the functional repertoire of bacteria central to human gut health.

## Main

The human gut microbiome is profoundly shaped by bacteriophages (phages), which regulate bacterial populations through lytic predation or temperate integration as prophages, phage–plasmids, and persistent virulent forms^1–3^. While lytic phages drive rapid bacterial turnover, temperate phages may promote longer-term coexistence through mechanisms such as superinfection immunity and lysogenic conversion, whereby prophages alter host phenotypes^4^. Together, these interactions facilitate extensive genetic exchange with other co-occurring phages^5^ and bacterial chromosomes^6^, contributing to the diversity and strong inter-individual variation that characterises the human gut virome^7^.

Despite this variability, a small number of phage lineages are consistently found worldwide^8–11^. However, both theoretical^12^ and observational^13^ studies indicate that prophages, which constitute the majority of gut-associated viruses, often persist in low-activity states. Their limited virulence and compact genomes, together with biases in current bioinformatic approaches, may constrain detection of low-abundance viral populations. As a result, efforts to catalogue the gut virome have largely focused on abundant phage genomes^11,14–16^, leaving the identity, biogeography, and ecological roles of globally distributed yet low-abundant bacteriophages unresolved.

Most currently known widespread phages infect *Bacteroidota* hosts^9,14,17–19^, even though bacteria from equally prevalent phylum *Bacillota* (formerly Firmicutes) are extensively infected by bacteriophages^11,18^. This disparity suggests that *Bacillota*-infecting phages can be systematically underdetected in current metagenomic analyses. Of particular interest are prophages integrated within bacterial lineages that have codiversified with human populations^20^, potentially disseminating globally alongside their hosts and superhosts. Recently, one such phage genus, *Mushuvirus* which infects *Faecalibacterium* and other short-chain fatty acid (SCFA)-producing *Bacillota* was identified in 1,300-year-old palaeofeces, with a genome sequence over 97% identical to modern references, indicating a long-term mutualistic relationship^21^. Despite displaying the hallmarks of a cosmopolitan lineage, including active replication^22^, genomic conservation, and a stable commensal host range^21^, *Mushuvirus* has not previously been recognised as a widespread component of the human gut virome.

Here, we apply phage-inclusive taxonomic profiling to the global collection of metagenomes, and reveal that despite its low abundance, *Mushuvirus* is the most prevalent bacteriophage genus among thousands of samples from six continents, defining, alongside a suite of co-occurring lineages, a core human gut virome at the threshold of metagenomic detection.

## Results

### A distinct family of human gut bacteriophages

The longstanding association of the *Mushuvirus mushu* with health-related bacteria prompted a global investigation into the evolution and ecology of these bacteriophages. We identified 528 *Mushuvirus*-related complete phage genomes from five comprehensive genomic catalogues spanning viral and bacterial diversity (Methods; Supplementary Table 1; Extended Data Fig. 1). Phylogenetic reconstruction revealed two major clades forming a monophyletic lineage distinct from all other classified *Caudoviricetes* (Fig. 1a). This separation was independently supported by single-gene phylogenies (Supplementary Fig. 1), a proteomic tree (Supplementary Fig. S2-3), and a genome-wide gene-sharing network (Fig. 1b; Supplementary Table 2), all of which supported a distant relationship to transposable Mu-like phages. On this basis, we propose designating the family Candidatus *Mushuviridae* as a distinct viral lineage, with two subfamilies, *Mulanvirinae* and *Lionvirinae*, characterised by different genomic and host-range features. The *Mushuviridae* pangenome shows a highly conserved modular organisation, typically encoding 48–60 proteins arranged into four functional modules: (1) integration, excision, and transcriptional regulation, (2) virion structure, (3) tail assembly, and (4) an accessory gene module undergoing targeted hypermutation by a DGR system in 9 out of 13 genera (Fig. 1c). Contrary to pervasive genomic mosaicism, typical of many phage families^23,24^, *Mushuviridae* showed higher sequence conservation than other prevalent human gut phage families (Fig. 1d).

**Fig. 1.**
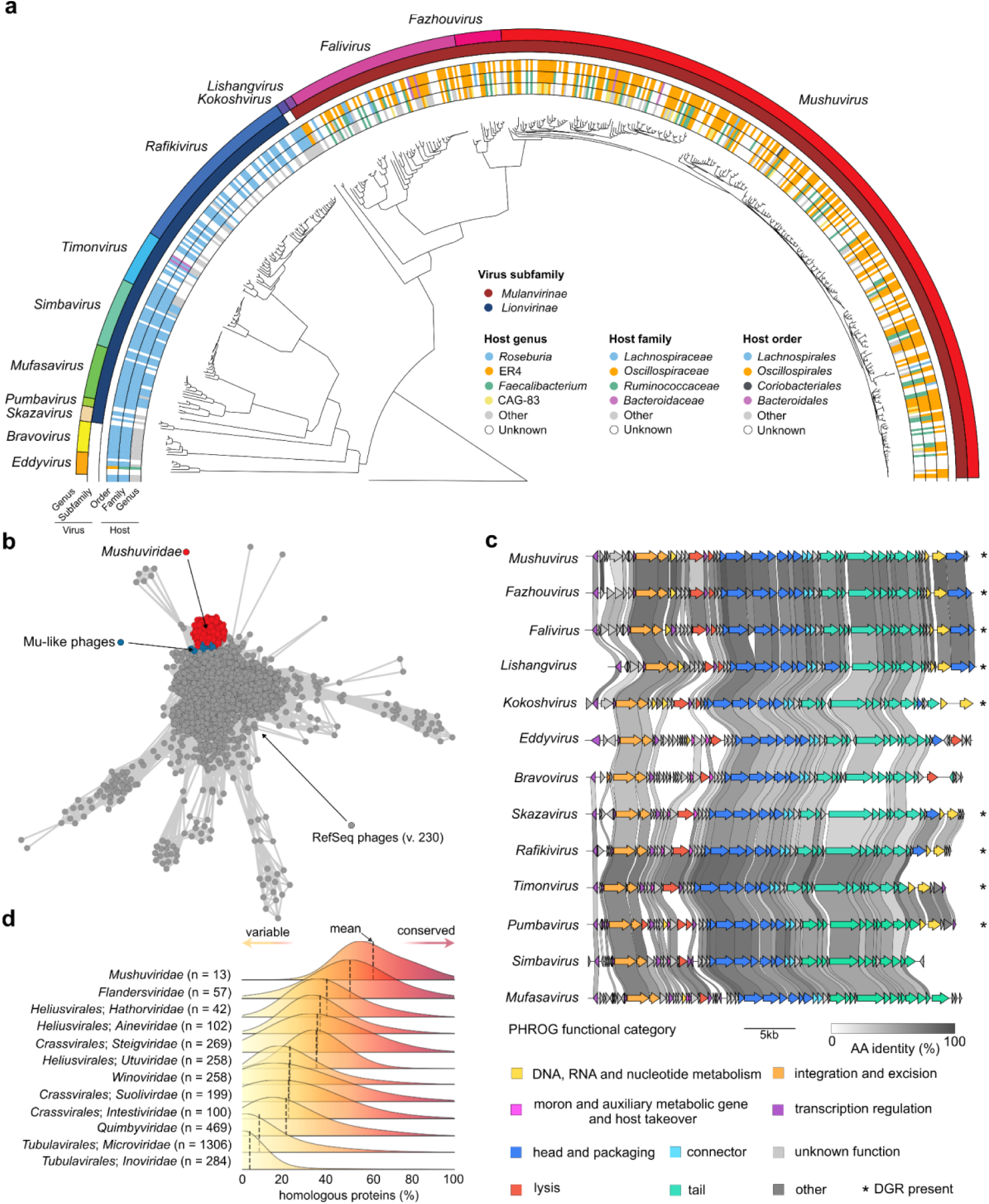
Phylogeny and conserved genome architecture of *Mushuviridae* phages that interact with health-associated gut bacteria. a, Proteomic tree (ViPTree) representing intragenomic diversity of candidate *Mushuviridae* genera. Outer-to-inner rings indicate the proposed viral genus and subfamily, as well as the host order, family, and genus as determined by prophage flanking regions (Extended Data Fig. 2). Assignments to the *Lachnospirales* and *Oscillospirales* orders largely align with the tree topology. The tree is rooted using the closest outgroup to *Mushuvirus*-related transposable phages (TBP_9153) integrated in *Enterocloster lavalensis*, selected via an approximate maximum-likelihood tree of terminase large subunit (TerL) genes (Extended Data Fig. 1c). **b,** Gene-sharing network (vContact3) showing the clustering of 528 *Mushuviridae* genomes relative to Mu-like phages from RefSeq (v230). **c,** Synteny in 13 representative *Mushuviridae* genomes (Clinker), showing the highly conserved four-module architecture. Asterisks indicate the presence of diversity-generating retroelement (DGR) systems within the accessory module. **d,** Comparative distribution of genomic conservation across prevalent human gut phage families. Values indicate the percentage of shared homologous proteins (MMseqs2; ≥30% sequence identity, ≥30% coverage, and *e*-value ≤10^-3^) between viral genera within each family. *Mushuviridae* (*n* = 13 genera) show the highest mean genomic conservation (dashed lines) compared to other major gut viral families.

By identifying prophage-flanking host genomic regions and using a lowest common ancestor approach (Methods), we assigned 265 of 528 *Mushuviridae* genomes (50%) to 87 bacterial species from 42 genera, 209 (40%) genomes to higher taxonomic ranks than species and 32 genomes were unassigned (Extended Data Fig. 2, Supplementary Table 3). In cases where left and right flanks were available, host associations were consistent in 98% of cases (391/398, Supplementary Table 4), and the remainder were binned into MAGs or found in bacterial isolates. Detected hosts were dominated by the orders *Lachnospirales* and *Oscillospirales*, largely, but not exclusively corresponding to two major clades in the *Mushuviridae* phylogeny (Fig. 1a). All assigned hosts encoded at least one metabolic pathway for the production of SCFA (Supplementary Fig. 4; Supplementary Table 5). The hosts include key taxa associated with human health, such as *Faecalibacterium*^25^, *Roseburia*^26^, *Blautia*^27^, *Dysosmobacter*^28^, CAG-170^29^ and *Enterocloster*^30^. The inferred host range varied widely between *Mushuviridae* genera, from narrow-host-ranges as observed for *Skazavirus* (infecting *Blautia*) and *Bravovirus* (infecting *Enterocloster*), to an exceptionally broad host range of *Mushuvirus* phages infecting up to 47 species across 21 genera (Extended Data Fig. 2). Collectively, these results provide detailed characterisation of a previously unknown viral lineage that maintains multi-host interactions with core members of the human gut microbiota.

### The most prevalent among human gut phages

To investigate the global distribution of the *Mushuviridae*, we used a highly sensitive taxonomic profiling approach (sylph^31^; see Methods) validated on simulated data to ensure accurate detection of viral genomes at low coverage (≥ 0.3) without misclassification (Supplementary Fig. 5). Using this approach, we identified *Mushuviridae* sequences in 89% of 53,976 human gut metagenomes worldwide (Fig. 2a; see Methods), in 99.5% crossing validated detection threshold, that positioned this previously undescribed lineage among the most widespread families *Winoviridae* (91%) and *Microviridae* (90%), as well as orders *Heliusvirales* (95%) and *Crassvirales* (89%). This ubiquity extends to the ancient human gut. We detected *Mushuviridae* in paleofeces from five archaeological sites in North America and Europe (Fig. 2b)^32–36^, including reconstruction of the *Mushuvirus mushu* genome (80% breadth of coverage) in the gut of the 5,300-year-old Tyrolean Iceman mummy (Fig. 2c).

**Fig. 2.**
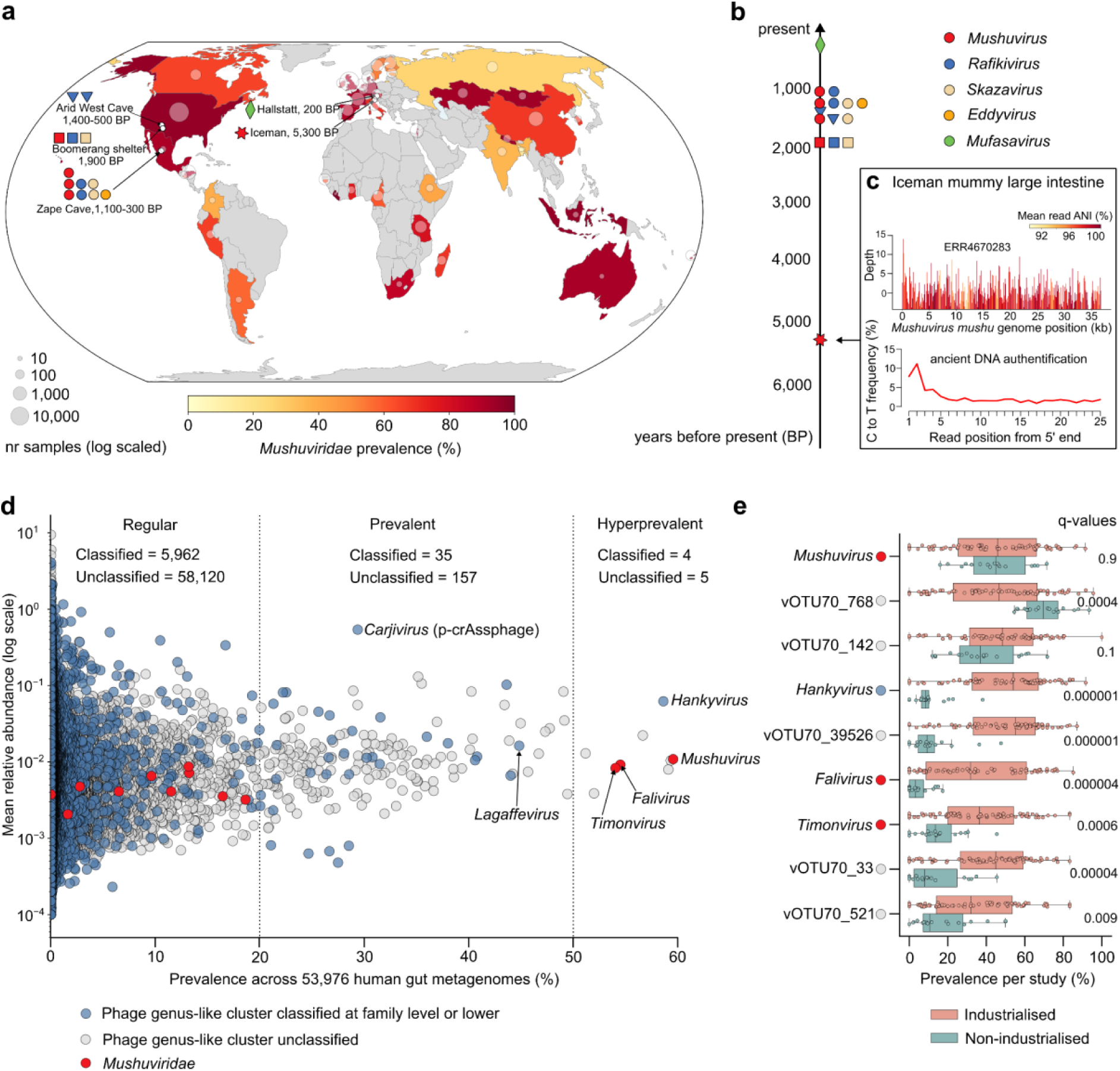
***Mushuviridae* phages are widespread and longstanding components of the human gut. a,** Global map of *Mushuviridae* prevalence calculated per country using sylph. Given uneven sampling indicated by log-scaled bubble sizes, observable patterns should be interpreted with caution. White dots indicate the geographical locations of archaeological sites/specimens where *Mushuviridae* were identified. **b,** Timeline presenting the widespread presence of *Mushuviridae* over time. **c,** Coverage and deamination plot confirming the reconstruction of an authenticated ancient *Mushuvirus mushu* genome from the gastrointestinal tract of the Iceman mummy Ötzi (ERR4670283). **d,** Abundance-prevalence plot showing distribution of genus-like clusters (70% ANI and 85% coverage) across 53,976 human gut metagenomes. **e,** Prevalence of nine hyperprevalent (>50%) phages in non-industrialised (n=18) and industrialised (n=79) studies. Statistical significance (q-values) was calculated using a two-sided Mann–Whitney U test with Benjamini–Hochberg FDR correction.

The *Mushuviridae* family’s global footprint is driven by three candidate genera (*Timonvirus*, *Falivirus*, and *Mushuvirus*), which belong to the group of 195 highly prevalent genera (the top 0.3% of 64,274 known gut viral genera) present in >20% of global samples at all continents, except Antarctica (Fig. 2d, Supplementary Table 6). Among them, *Mushuvirus* is the most cosmopolitan, detected in 60% of samples across both industrialised and non-industrialised populations (Fig. 2e). This homogeneous global distribution contrasts with other prevalent gut viruses, such as the *Bacteroidota*-associated *Carjvirus* (p-crAssphage) and *Hankyvirus*, whose prevalence typically declines in non-industrialised settings (from 32% to 11%, and 64% to 10%, respectively). While human superhost lifestyle did not influence *Mushuvirus* carriage, age emerged as an important factor, with prevalence increasing from 1% in newborns (n = 2,672) to 56% in adults (n = 10,651), indicating continuous colonisation throughout childhood (Supplementary Fig. 6). Furthermore, *Mushuvirus* prevalence declined from 55.2% in healthy individuals (n = 580) to 29.5% in patients with inflammatory bowel disease (IBD; n = 868). IBD samples showed a significant increase in relative abundance (q = 0.006, Extended Data Fig. 3), consistent with previous findings linking inflammation with higher temperate phage activity^22^.

We next sought to understand why *Mushuvirus* had remained undetected in thousands of metagenomes worldwide despite its high prevalence. First, *Mushuvirus* is consistently maintained at very low relative abundances (range: 0.0001% to 0.7142%; mean = 0.01% ± 0.02%), even in longitudinally sampled, viral-enriched datasets, where it remains active (Supplementary Fig. 7). Second, as mushuviruses are commonly integrated into their bacterial host genome, the reference sequence in the database may be represented as a phage-bacterial chimera. To test this hypothesis, we investigated the *Mushuvirus* representative sequence (vOTU-007404) in the Unified Human Gastrointestinal Virome catalogue (UHGV)^16^, which aggregates viral genomes published in 12 previous studies. Interestingly, vOTU-007404 is a 64 kb sequence, whereas the *Mushuvirus* genome is 36 kb. Further investigation revealed that this sequence is a phage–phage chimera (Supplementary Fig. 8), that might reflect a tandem integration event or represent a misassembly. This leads to problems in identifying such sequences as contamination using automatic quality control methods (CheckV^37^) which predict all encoded proteins as viral and classify the sequence as a complete genome. Combined with the stringent identification thresholds (75% horizontal coverage)^38^ used in previous large-scale studies^16,39^, these biases may have contributed to missing *Mushuvirus* as a major component of the human gut virome.

### Phylogeography and codiversification with humans

Phylogenomic analysis of the *Mushuvirus* genus revealed a geographically restricted subpopulation centered in Europe, North America, and Western and Central Asia, with notably sparse representation in East Asia (Fig. 3a). Based on recommendations of International Committee on Taxonomy of Viruses (ICTV) to classify phage species as sharing ≥95% nucleotide identity over their full genome length^40^, we proposed the delineation of this lineage as a separate species, *Mushuvirus draco* (Supplementary Fig. 9; Supplementary Table 7). Despite its wider distribution, *Mushuvirus mushu* genomes were slightly more conserved (mean intra-species tANI = 96.6% ± 1.2%) than the geographically restricted *Mushuvirus draco* (95.7% ± 1.4%). Population genetic analyses identified three 500–600 nt regions of high nucleotide variability (Fig. 3b; Supplementary Table 8); however, none of these overlapped with region of high genetic differentiation between the two *Mushuvirus* species (mean F_ST_ = 0.79 ± 0.06; Fig. 3c). In *Mushuvirus mushu*, this locus encodes two to three small hypothetical genes, whereas *Mushuvirus draco* and other *Mushuviridae* (*Fazhouvirus*, *Falivirus*, and *Lishangvirus*; Fig. 1c) carried a single gene. This gene was annotated based on sequence and structural homology as an alpha-amylase, chitinase, or carbohydrate-binding protein.

**Fig. 3.**
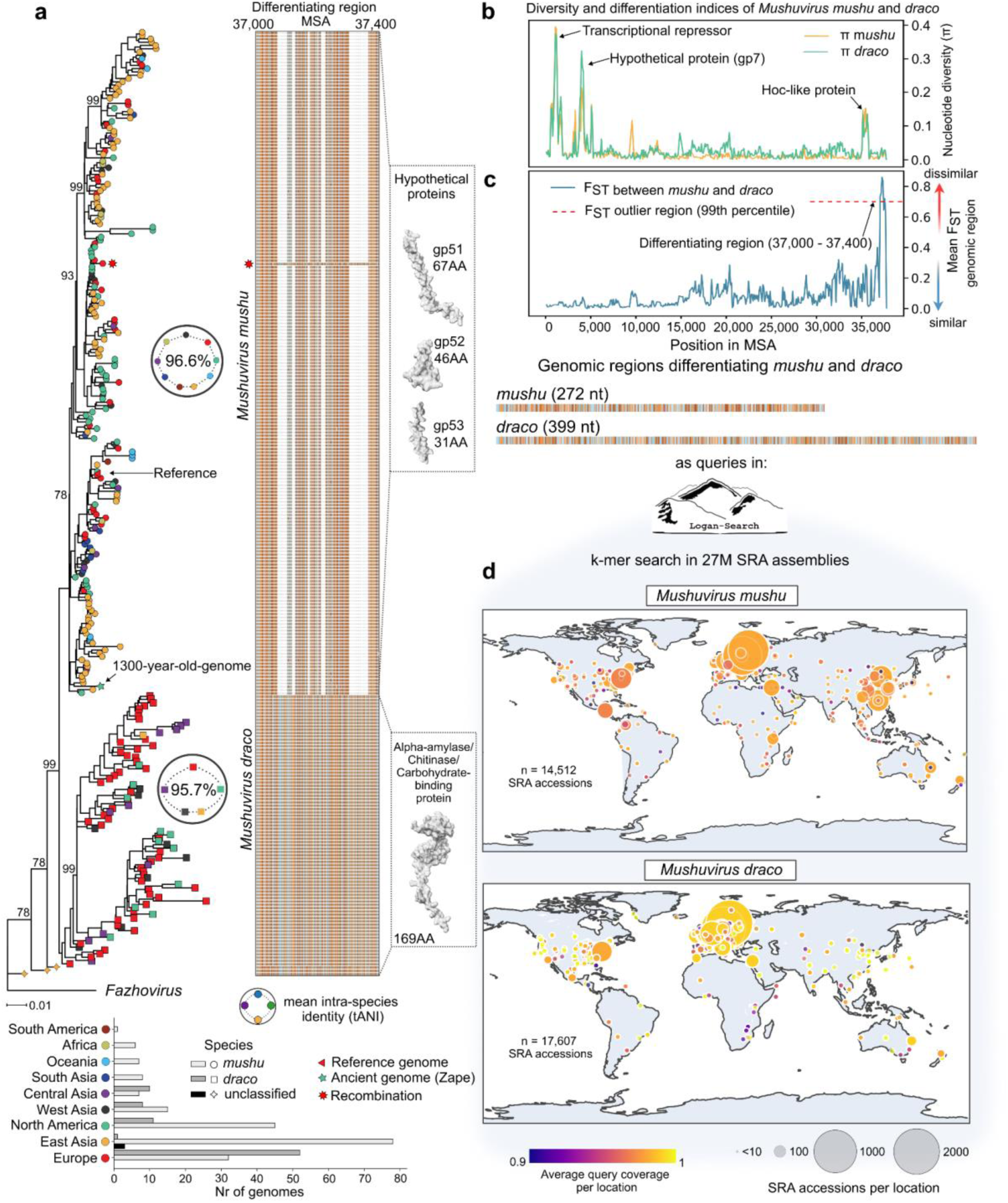
Population genomics of *Mushuvirus* reveal cryptic species dominating in Western countries. a, Maximum-likelihood phylogenetic tree of 284 *Mushuvirus* genomes, rooted with *Fazhovirus*. The phylogeny includes the reference genome from France (NC_047913.1, red triangle) and the 1300-year-old genome from Zape Cave, Mexico (green star)^21^. Tips are colored by geographic origin. Insets show mean intra-species identity (tANI) and a multiple sequence alignment of the differentiating region (positions 37,000-37,400) demarcating *Mushuvirus mushu* and *Mushuvirus draco* species. The red star by the alignment indicates a single inter-species recombination event of the region from *Mushuvirus draco* into *Mushuvirus mushu*. Protein structures for the differentiating genes were predicted by AlphaFold 3 and manually annotated via BLASTp (NCBI ClusteredNR database) and structural homology (Foldseek). AA: amino acids. **b,** Nucleotide diversity (π) and **c,** genetic differentiation (fixation index, F_ST_) calculated in 100-bp sliding windows across the *Mushuvirus* genomes. Three diversity peaks correspond to a transcription repressor, hypothetical protein gp7, and a hoc-like protein. Red points indicate F_ST_ outliers (99^th^ percentile). **d,** Sequences of differentiating region between *Mushuvirus mushu* and *Mushuvirus draco*. **e,** Global geographic distribution of species-specific *k*-mers identified in the Logan database (representing assemblies from 27 million SRA accessions). Circles are scaled by the number of SRA accessions per location and colored by average query coverage of the differentiating region.

To resolve the global biogeography of both *Mushuvirus* species, we queried the Logan database with assemblies from 27 million Sequence Read Archive (SRA) accessions^41^ using two sequences from the differentiating region (Fig. 3d), and identifying their presence in 32,119 SRA samples (Supplementary Table 9). This large-scale census confirmed the geographical partition observed in the phylogeny (Fig. 3a), with *Mushuvirus mushu* being near universally ubiquitous, and *Mushuvirus draco* showing marked dominance in Europe and the United States (Fig. 3e). To elucidate whether this speciation reflects simple geographical separation or a specific association to human lineages, we combined our taxonomic profiles with mitochondrial DNA haplogroups recovered from publicly available human gut metagenomes of healthy individuals (Supplementary Table 10)^42^. *Mushuvirus draco* and *Falivirus*, which both encode specific carbohydrate-binding proteins in the differentiating region, are more prevalent in individuals with the West Eurasian haplogroups R, U, H, T, X, and less prevalent in African (L) and East Asian, Siberian, and Native American (A, B, C, D, F) haplogroups (Supplementary Fig. 10a,b). This association is preserved in different geographical regions, as individuals in both Canada and Israel who carry the most common West Eurasian haplogroup (H)^43^ show enrichment of *Falivirus* relative to other people from these countries (Supplementary Fig. 10c). This analysis suggests that the distribution of these dominant gut phages is not merely a product of geographical separation but may reflect the deep tripartite relationship with host bacteria^20^ and human superhosts that codiversify together.

### Prophages expand the phenotype of Faecalibacteria

Given the widespread occurrence of *Mushuvirus* phages and their association with SCFA-producing hosts, we examined their functional contribution to the health-promoting butyrate-producer *Faecalibacterium duncaniae* A2-165 (formerly *F*. *prausnitzii*)^25^. Transcriptomic profiling identified five prophage-encoded genes expressed above the host bacterial average (Fig. 4a; Supplementary Table 11)^44^, which were independently confirmed via LC–MS/MS proteomics (Fig. 4b; Supplementary Table 12)^45^. Three proteins, including a putative repressor, a major capsid protein, and a Hoc-like receptor-binding protein were translated in the whole–bacterial culture, with the latter also present in the extracellular vesicle (EV) fraction (Fig. 4c, Supplementary Table 13), indicating active export beyond the bacterial cell.

**Fig. 4.**
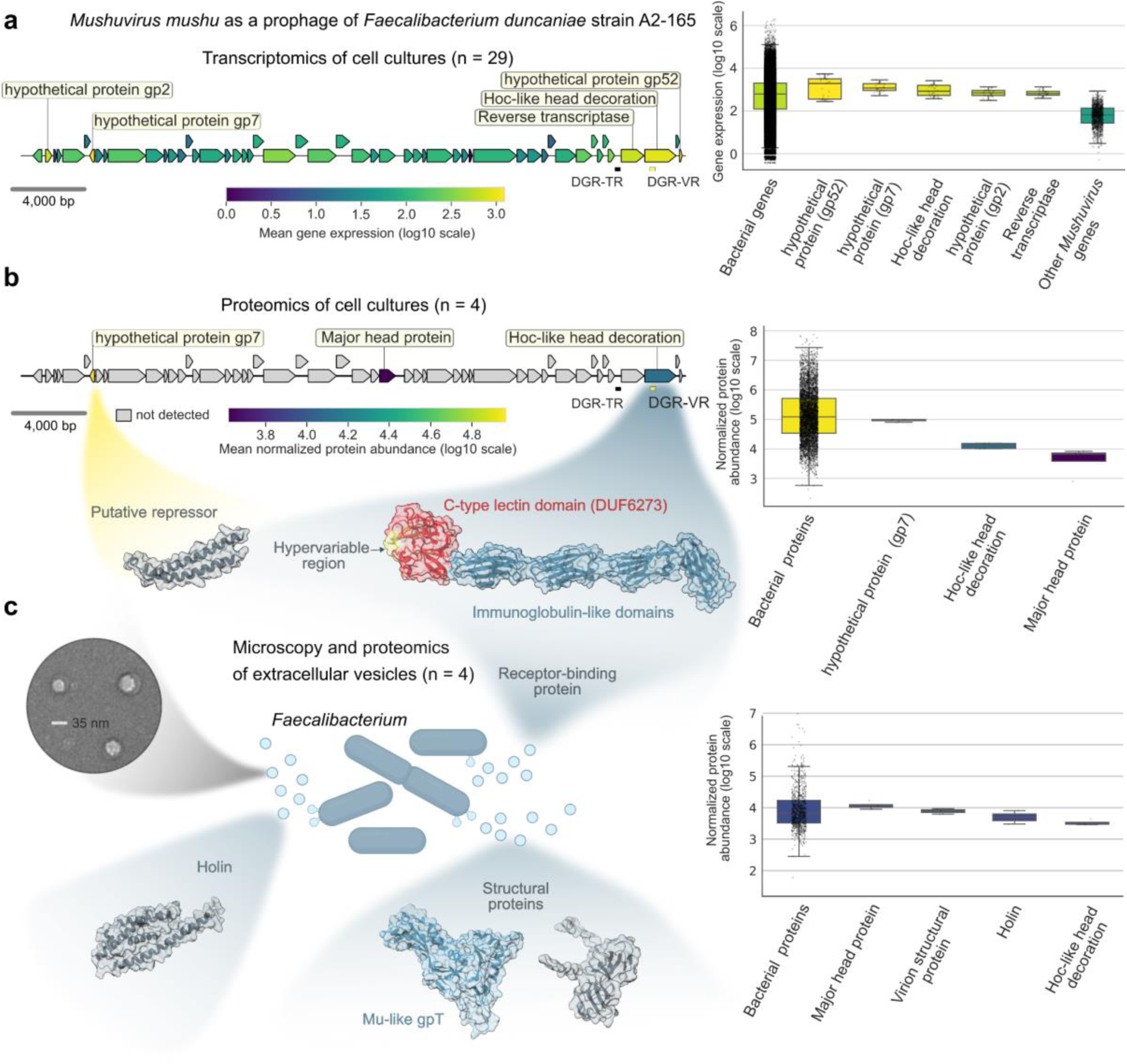
***Faecalibacterium* bacteria express and secrete *Mushuvirus*-encoded proteins. a,** Summary genome plot showing the transcriptomics results based on three RNA-seq datasets from *Faecalibacterium duncaniae* (formerly *F*. *prausnitzii*) grown under different conditions (see: Methods). Five prophage-encoded genes are expressed at levels higher than the mean value for bacterial genes (433 ± 9.84). **b,** Results of a *Faecalibacterium duncaniae* proteomic analysis using data generated independently of the transcriptomic datasets. Hypothetical protein 2 and Hoc-like protein are consistently transcribed and translated, contributing to the *Faecalibacterium* functional repertoire. **c,** Results of proteomic analysis of *Faecalibacterium duncaniae* extracellular vesicles reveal the presence of three functional categories of prophage-encoded proteins. Active expression (panel a and b) with concurrent presence of phage-encoded receptor-binding proteins in the bacterial secretome, lacking phage particles in electron microscopy (left; scale bar: 35 nm), strongly suggests a functional role of this protein for the bacterial host.

The Hoc-like protein represents the tentaclin family^46^, which contains immunoglobulin-like domains and a C-type lectin domain (DUF6273). It has a DGR-targeted hypervariable region (VR) that likely mediates interactions with superhost or microbial carbohydrates. Phylogenetic analysis of this locus showed that it is almost completely different from the overall tree (Robinson–Foulds distance: 0.95; Supplementary Figure 11). It resolved four major variants, structured by geography rather than host taxonomy, with distinct clusters dominating in East Asian and European populations (Extended Data Fig. 4a; Supplementary Figure 11). Sequence variation was concentrated within the ∼30-amino acid VR segment of the lectin domain, contrasting sharply with the high conservation observed across the rest of the genomes (Extended Data Fig. 4b).

This localised hypervariability differs from the host-specific VR variation observed in the broad-host-range *Lagaffevirus* prophage (also integrated in *F. duncaniae*) and instead resembles patterns recognised in hankyviruses (Extended Data Fig. 4c and 4d). This extensive variation is driven by a highly efficient DGR system that remains active during host lysogeny (Fig. 4e), resulting in the high sequence variation that we observed in spontaneously produced viral particles from *F. duncaniae* cultures (Extended Data Fig. 4e). This provides a continuous reservoir of phages with diverse binding specificities.

Together, our results demonstrate that a prophage-encoded receptor-binding protein is acquired by bacterial host strains, actively diversified via DGR, and secreted into the gut environment for interactions with carbohydrates present on human or bacterial cells. This mechanism represents a clear example of phenotype extension through lysogenic conversion of bacterial hosts.

## Discussion

More than two decades^47–49^ of metagenomic studies on the human gut microbiome and virome have generated a vast reservoir of sequence data, transforming our understanding of phage diversity and their interactions with bacterial hosts^13,16^. Advances in sequencing and computational approaches have enabled recovery of complete viral genomes from complex communities, revealing globally distributed phage lineages, including crAssphages^14^, and other previously uncharacterized groups^10,17,18^. Despite this progress, the gut virome remains largely unresolved, with hundreds of predicted viral families lacking formal taxonomic classification^16^. This lack of resolution hinders cross-study comparability, masks reproducible associations with health and disease, and complicates the prioritisation of phages for mechanistic studies. Here, we address this gap by characterising the *Mushuviridae*, a previously unrecognised yet highly prevalent viral family. Additionally, our global-scale profiling of gut phages defines a core virome that is consistently detected in diverse human populations, and identifies cosmopolitan, but uncharacterized viral lineages for future studies.

Previous observations indicate that the human gut virome is highly individual-specific and stable^7^, with limited distribution of low-rank taxonomic groups^10^. Our observation that 99.7% of phage genera occur in <20% of global samples corroborates this. This raises a key question: what evolutionary adaptations enable the remaining 0.3% of lineages to achieve global prevalence? Broad host range is considered a primary driver of phage distribution, facilitated by DGR systems and DNA methyltransferases^16^. However, these features alone did not explain the prevalence patterns observed for *Mushuviridae* genera. An alternative, and potentially complementary, mechanism is phage adherence to intestinal epithelial surfaces. Recent work has linked such adherence to increased prevalence in human gut metagenomes^50^, and *Mushuviridae* may achieve this through their frequently encoded carbohydrate-binding proteins, as described by the bacteriophage adherence to mucus (BAM) model^51^ whereby lytic phages provide a non–host-derived antimicrobial defence by evolving enhanced persistence to a mucosal surface^52^. Although *Mushuviridae* are temperate phages and the carbohydrate-binding proteins are expressed by the bacterial host, the principle of enhanced retention within the gut environment may be shared. Together, these findings suggest that phage–host interactions in the gut should be considered not only at the level of bacterial hosts but also in the context of direct interactions with the human superhost. Such a perspective may help explain the striking contrast between rare and highly prevalent taxa, as well as the phylogeographic patterns observed in this study.

Although phages are traditionally viewed through the lens of predation, mutualistic interactions are increasingly recognised as drivers of bacterial trait evolution in complex communities^4^. The *Mushuvirus* genus offers an exemplary model of such a relationship; its coexistence with bacterial hosts is associated with a stable global presence in nearly 60% of humans and genome conservation over thousands of years. This persistence likely reflects a trade-off between low-level replication and genomic innovations provided by the prophage to the host. As we demonstrated in our multi-omics experiments with health-associated Faecalibacteria and *Mushuvirus* prophages, bacterial cells actively express a viral-encoded DGR system during lysogenic conversion that increases variability in receptor-binding proteins, which may interact with carbohydrates on bacterial cell walls and/or human epithelial or immune cells. We anticipate that future studies will further elucidate these tripartite interactions, potentially paving the way for the incorporation of specific phage lineages into microbiome-based medical applications.

## Methods

### Data collection for *Mushuviridae* identification

Six genomic resources were integrated to identify *Mushuvirus*-related sequences. The first was the Unified Human Gastrointestinal Virome (UHGV)^16^ v1.0, a comprehensive catalogue comprising 208,604 high-quality or complete viral genomes obtained from 12 human gut studies. The second dataset was IMG/VR^53^ v4.1, which contains 518,882 high-quality or complete viral genomes derived from diverse environments. The third resource comprised 9,766 complete transposable phage genomes extracted from RefSeq bacterial genomes (Transposable phages: TBP)^54^. The fourth was the Unified Human Gastrointestinal Genome (UHGG)^55^ v2.0.2, a collection of 289,232 MAGs from the human gut. The fifth dataset included two collections of non-human primate (NHP) gut metagenomes: one spanning 23 taxa^56^ and another consisting of baboon faecal metagenomes^57^. The sixth was a previously reported complete ancient *Mushuvirus* genome from a prior study (Genbank accession: BK063464.1)^21^.

### Data collection for phage-inclusive reference database

A comprehensive reference database for taxonomic profiling of the human gut virome was constructed by integrating viral genomes from multiple sources. These included 528 genomes representing 13 *Mushuviridae* genera, 6,741 ICTV-classified bacteriophages (ICTV’s Virus Metadata Resource, VMR v40.1), 1,032 genomes from the *Heliusvirales* clade^10^, 1,776 Gubaphages^18^, and 13,956 predicted prophages identified in UHGG and NCBI bacterial genomes^1^. Prophage candidates were retained only if classified as at least medium quality by CheckV^37^ v1.0.1 and predicted to be viral by geNomad^58^ v1.8.1. Also included were 134 inducible human gut prophages^19^, 30 *Flandersviridae*^59^, four loVEphages^17^, one *Hankyvirus*^9^, 97,809 medium, high-quality or complete genomes from UHGV^16^, and 4,744 prokaryotic-species representatives from UHGG^55^. All viral genomes were clustered into species-level groups using Vclust^60^ v1.3.1 at 95% average nucleotide identity (ANI) and 85% alignment fraction, resulting in 102,336 phage clusters (vOTU95). Cluster representatives were selected with priority given to genomes with established taxonomy and manually curated completeness (Supplementary Table 14).

### Metagenome collection for taxonomic profiling

For taxonomic profiling, 53,976 publicly available shotgun metagenomic samples from 90 studies of the human intestinal tract were used (Supplementary Table 15). Seventy-one studies were selected from CuratedMetagenomicData^61^ v3.21, with 19 additional studies included to increase geographic, demographic, and clinical representation, mostly collected through Metalog^62^. These samples span different age categories (2,672 newborns, 930 children, 10,651 adults, 1,280 seniors, and 37,911 with unreported age), genders (7,377 males, 7,933 females, and 38,666 with unreported gender), lifestyles (49,100 industrialised and 3,196 non-industrialised), 42 distinct health conditions, and 42 countries. Metadata for palaeofaeces and ancient gut content samples representing nine studies were retrieved from AncientMetagenomeDir^63^ v25.09 (Supplementary Table 16). For subsequent analyses, only metagenomes with >10M reads were retained.

### Identification of *Mushuviridae* sequences

*Mushuvirus*-related sequences were identified by searching public viral genome collections (see “Data collection for *Mushuviridae* identification” Methods section) using the reference *Mushuvirus mushu* TerL (AUV61537.1) with MMseqs2^64^ v13.45111 (options: easy-search --min-aln-len 590 -e 0.05 --max-seqs 10000 -s 5.0). Protein sequences with an identity ≥ 0.5 were aligned with FAMSA^65^ v2.2.3 and converted into an HMM profile using hmmbuild^66^ v3.3. This profile was used to search for distant TerL homologs using hmmsearch v3.3 (bit-score ≥ 50). The resulting sequences were realigned with FAMSA, and the alignment refined by removing positions with >80% gaps (trimAl^67^ v1.5, option -gt 0.2), and excluding sequences with >60% gap content.

An approximate maximum likelihood tree was inferred using VeryFastTree^68^ v3.1.0 and annotated with bit-score values in TreeViewer^69^ v2.2.0. Sequences of the monophyletic lineage with bit-scores ≥ 1000 were selected for downstream analyses. *Mushuvirus*-related contigs were clustered at 70% ANI and 85% alignment fraction of the shorter sequence using Vclust^60^. Sequences from each cluster were aligned with MAFFT^70^ v7.5.0.5 (options: --maxiterate 1000 --localpair) and manually inspected to identify conserved 12-bp integration motifs defining phage genome ends. These motifs were used to separate viral genomes from variable bacterial regions (a list of all motifs is provided in Supplementary Table 17), ensuring 100% complete bacteriophage genomes, which were then used in a BLASTn^71^ v2.16.0+ search for additional genomes integrated into MAGs from the UHGG and contigs from NHP.

### Demarcation of the *Mushuviridae* family

The internal genomic diversity of *Mushuviridae* lineage was visualized using ViPTreeGen^72^ v1.1.3 and GraPhlAn^73^ v1.1.3. Demarcation of this lineage as a separate family was supported by three independent lines of evidence. First, a phylogenetic framework was established using TerL protein; related ICTV-classified phages were identified from VMR v40.1 using HMM-based recruitment strategy described above (see: Identification of *Mushuviridae* sequences). Homologous proteins were aligned with FAMSA and maximum likelihood tree was inferred using IQ-TREE^74^ v2.1.4 (options: -nt AUTO -m MFP -B 1000 -alrt 1000 --runs 10). Second, *Mushuviridae* were integrated into a global proteomic tree of RefSeq v220 phage genomes using the VipTree v4 server (https://www.genome.jp/viptree/)^72^. Third, a gene-sharing network was constructed in vContact3^75^ v3.0.5 using an updated RefSeq v230 genome collection and visualised with Cytoscape^76^ v3.10.1.

Genomic characteristics of the family were determined by functional annotation of all *Mushuviridae* genomes using Phold^77^ v0.2.0, followed by synteny analysis of genus-level representative sequences using Clinker^78^ v0.0.31 via CAGECAT web service (https://cagecat.bioinformatics.nl/). Genetic heterogeneity across *Mushuviridae* and other prevalent phage families was measured using MMseqs2. Pairwise percentages of shared genes were calculated at the minimum threshold of detectable homology (≥30% sequence identity, ≥30% coverage, and *e*-value ≤10⁻³), using one representative genome sequence per genus within each studied family.

### Host assignment

Two complementary strategies were applied to assign bacterial hosts to *Mushuviridae* phages: extraction of flanking bacterial sequences from contigs and direct host inheritance from genomes identified as prophages. In the first approach, flanking bacterial sequence fragments (≥30 bp) were extracted by trimming contigs at the identified conserved 12-bp integration motifs and mapped against all UHGG genomes using BLASTn. Alignments with ≥99% query coverage were retained, and the host taxon was assigned based on the hit with the highest sequence identity. In cases where multiple hits met these criteria, a lowest common ancestor (LCA) approach was applied to determine the host lineage. In the second approach, host taxonomy was derived directly from the source genomes, either UHGG MAGs or Refseq bacterial isolate genomes (TBP), in which *Mushuviridae* sequences were identified as integrated prophages.

### Taxonomic profiling

To overcome the limitations of assembly-based metagenomics, which often fails to recover genomes from low-abundance populations, we used sylph^31^ v0.9.0, a high-sensitivity taxonomic profiler that employs *k*-mer-based containment sketches to detect and quantify genomes at low sequencing depth. The performance of this tool for detecting *Mushuviridae* was validated using 100 simulated metagenomes, generated using InSilicoSeq^79^ v2.0.1. Each simulation contained equal representation of all *Mushuviridae* genera, with per-genus coverage ranging from 0.1× to 10×, a mean read fragment length of 100 bp (standard deviation = 1), and no additional bacterial or phage sequences. These metagenomes were analysed with sylph using a reference database of 102,336 vOTU95 and 4,744 UHGG species (see: Data collection for phage-inclusive reference database), constructed with a compression parameter optimised for low-coverage genomes (-c 100). This validation established the minimum coverage threshold required for reliable detection and quantified misclassification rates against non-*Mushuviridae* references (Supplementary Fig. 5).

Global-scale, phage-inclusive profiling of human gut metagenomes was performed using a customised pipeline. Specifically, publicly available metagenomes (see: Metagenome collection for taxonomic profiling) were downloaded using fastq-dl v3.0.1, and sequencing adapters and low-quality bases were removed using Cutadapt^80^ v5.1 via Trim Galore^81^ v0.6.10. Quality-controlled reads were sketched and profiled using sylph (-c 100 -t 50 --min-number-kmers 30 -u). The reference database was used at the species level, with genomes further aggregated into genus-level groups (vOTU70: 70% ANI and 85% coverage) using Vclust (Supplementary Table. 14). Prevalence and abundance metrics were summed per vOTU70. Manual curation of the fifteen most prevalent vOTU70 identified four phage-bacterial chimeric genomes that were excluded from downstream analyses (Supplementary Fig. 12).

### Ancient DNA analysis

Paleofecal and ancient gut metagenomes were screened using the taxonomic profiling pipeline described for modern samples. Because ancient, highly degraded reads might be potentially missed by sylph, in the oldest sample (derived from the Iceman mummy large intestine), trimmed reads were additionaly mapped to *Mushuvirus mushu* reference genome (NC_047913.1) using Bowtie2^82^ v2.4.1 with parameters modified to improve the recovery of degraded DNA reads by allowing for one mismatch in the seed (--very-sensitive, -N 1, -L 20, --no-unal). Mapped reads were used to authenticate the ancient origin of the viral sequences by estimating DNA deamination patterns in MapDamage2^83^ v2.2.3

### Associations with IBD

Association of *Mushuviridae* genera with IBD were evaluated using compiled shotgun metagenomic samples from healthy individuals and IBD patients across five studies^84–88^. Per-study odds ratios (ORs) were computed for each genus independently and summarised via a random-effects meta-analysis using the DerSimonian–Laird approach. Within-study variances were derived from 2×2 contingency tables using Haldane–Anscombe correction (adding 0.5 to all cells to prevent undefined odds ratios and stablize variance estimates). The statistical significance of pooled estimates was assessed using two-sided Z-tests. For individual studies, the significance of ORs was assessed by permutation testing (10,000 permutations), with condition labels shuffled without replacement. The Benjamini–Hochberg false discovery rate (FDR) correction was applied independently to each analysis, with significance defined as FDR < 5%.

Differences in relative abundance and genome coverage (per 1M reads) between IBD and healthy groups were assessed using two-sided Mann–Whitney U tests with Benjamini–Hochberg FDR correction applied across all genera. Relative abundance (sequence_abundance) and coverage (True_cov) were extracted from taxonomic profiles generated by Sylph. Genus-level prevalence was calculated separately for IBD and control groups as the proportion of samples in which each genus was detected. All statistical analyses were performed in Python using SciPy^89^ v1.11.4 and statsmodels v0.14.1.

### Phylogeographic and population genomic analysis

The phylogeography of *Mushuvirus* was reconstructed using all identified complete *Mushuvirus* genomes and the representative genome of *Fazhouvirus* as an outgroup. A whole-genome MSA was calculated with MAFFT^70^ (options: --maxiterate 1000 --localpair), and a maximum likelihood tree was inferred with IQ-TREE^74^ (options: -nt AUTO -m MFP -B 1000 -alrt 1000 --runs 10). The tree was annotated with metadata using the ggtree^90^ v3.16.3 package in R.

Genome-wide nucleotide diversity (π) and fixation index (F_ST_) were estimated from the MSA to characterise the evolutionary pressures differentiating *Mushuvirus* species. For each species, π was calculated using a custom Python script (Biopython^91^ v1.81 and NumPy^92^ v1.26.4) where per-site diversity was defined as the mean pairwise difference among valid nucleotides (A, T, G, C), excluding ambiguous bases and gaps. Diversity values were averaged in non-overlapping 100-nucleotide windows to generate genome-wide profiles. Genetic differentation between *Mushuvirus mushu* and *Mushuvirus draco* was quantified using Hudson’s F_ST_ estimator via scikit-allel v1.3.1. The MSA was converted to a haploid genotype matrix, with invariant or missing sites excluded, and allele counts computed for each species. Per-site F_ST_ values were averaged across 100-nucleotide windows; those in the top 1% (99th percentile) were classified as outliers, potentially representing regions under natural selection differentiating the two *Mushuvirus* species.

### Global distribution and human ancestry association

Sequences from the differentiating region between *Mushuvirus* species were used as queries for *k*-mer-based searches (*k* = 31) across all SRA assemblies (as of December 2023) in the Logan database^41^. Global distribution maps of *Mushuvirus mushu* and *Mushuvirus draco* distribution were generated based on Logan-search output. To investigate associations with human ancestry, mtDNA haplogroups, imputed from metagenomes in an independent publication^42^, were linked to our taxonomic profiles based on shared sample identifiers. *Mushuviridae* presence was extracted from respective taxonomic profiles, and prevalence was calculated for each major haplogroup (*n* >10 samples) and visualised as heatmaps. Due to high overall genomic diversity between *Mushuvirus mushu* and *Mushuvirus draco*, their presence in different haplogroups was determined using the Logan-derived occupancy data.

### Transcriptomics and proteomics

Transcriptomic data for *Faecalibacterium duncaniae* strain A2-165 (GenBank accession: CP022479.1) were derived from 29 samples spanning three independent studies^93–95^ uniformly analysed by Auger et al., 2022^44^. Briefly, preprocessed reads were mapped to the reference genome and quantified at the transcript level using HTSeq-count^96^ v0.12.4. The resulting count table was integrated with *Mushuvirus* gene annotations via shared locus tags (Supplementary Table 11) and expression values were log-transformed for visualisation. All details on transcriptomic profiling can be found in the original publication^44^.

Proteomics analyses were based on four publicly available samples representing the whole *Faecalibacterium duncaniae* culture^45^ and four newly generated data from EVs fraction extracted in independent experiment. Proteomics analyses of EVs fraction was performed by short SDS-PAGE followed by in-gel tryptic digestion and LC-MS/MS analysis on a timsTOF Pro mass spectrometer (Bruker) in dia-PASEF mode, with protein identification carried out using DIA-NN^97^ v1.8 in library-free mode against the *Faecalibacterium duncaniae* reference proteome at 1% FDR. Relative quantification was performed using MCQR^98^ v1.0.2, with protein intensities computed by summing normalized intensities of repeatable, protein-specific peptides. All details on proteomic profiling of bacterial cultures can be found in the original publication^45^ and for the EVs in the Supplementary Methods.

### Analysis of Hoc-like proteins and DGR-VR activity

A dendrogram showing the sequence relationships of *Mushuvirus* Hoc-like genes was generated with ggdendro v0.2.0 using a pairwise matrix of *Mushuvirus* Hoc-like gene sequence calculated with Vclust. MSAs were generated using MAFFT^70^ (options: --maxiterate 1000 --localpair) and merged with the dendrogram using ggtree^90^. DGR-VRs were predicted for each phage genus using the MyDGR^99^ webservice (https://omics.informatics.indiana.edu/myDGR/). Variable amino acid regions were extracted from alignment calculated with FAMSA^65^, and global pairwise similarities of DGR-VR regions were calculated using a Needleman-Wunsch algorithm in the PairwiseAligner module in Biopython^91^.

The activity of DGR-VR in *Mushuvirus* and *Lagaffevirus* prophages was assessed by mapping DNA reads from the supernatant of *Faecalibacterium duncaniae* strain A2-165 cultures to the reference genome (CP022479.1) using Bowtie2^82^ (parameters: -N 1 -L 32). Cultural conditions and sequencing library preparation protocols are described in detail in the original study^22^.

## Data availability

Databases included in this study: UHGV v1.0 (https://uhgv.jgi.doe.gov/), IMG/VR v4.1 (https://genome.jgi.doe.gov/portal/IMG_VR/), TBP (https://data.cyverse.org/dav-anon/iplant/home/zhangmujie/TBPGD/TBPGD.zip), NHP (https://zenodo.org/records/4641870; MGnify: MGYS00006808), and ancient *Mushuvirus mushu* genome (NCBI accession BK063464). Accession codes for metagenomes used in this study are available in Supplemenetary Table 15. Raw EV proteomics data have been deposited in the PRIDE repository under accession code PXD073554. All *Mushuviridae* and reference genome sequences used to construct the sylph database, as well as the resulting taxonomic profiles and other supplementary data, are available in Zenodo (https://doi.org/10.5281/zenodo.20084670).

## Code availability

Pipeline developed for phage-inclusive taxonomic profiling of human gut metagenomes is publicly available at GitHub (https://github.com/rozwalak/Mushuviridae_profiling)

## Acknowledgments

We thank Jeffrey K. Cornuault and Marianne De Paepe, who discovered and described the first *Mushuvirus mushu*, for helpful discussions and for sharing unpublished data with us. This work was supported by NWO, the European Research Council (ERC) Consolidator grant 865694: DiversiPHI, and the Deutsche Forschungsgemeinschaft (DFG, German Research Foundation) under Germany’s Excellence Strategy – EXC 2051 – Project-ID 390713860, the Alexander von Humboldt Foundation in the context of an Alexander von Humboldt-Professorship founded by the German Federal Ministry of Education and Research, grant (ID-AAF) from the Carnot Institute Qualiment© with funding provided by the French National Research Agency, and the program ‘Perły Nauki’, project number PN/01/0063/2022 (to P.R.) funded by Polish Ministry of Science and Higher Education. This work has benefited from the facilities and expertise of the Plateforme Analyse Protéomique de Paris Sud-Ouest (PAPPSO) which is supported by INRAE, the Ile-de-France regional council, IBiSA, and CNRS, and the Plateforme de Microscopie et Imagerie (MIMA2). P.R. was supported by Add-on-Fellowship from the Joachim Herz Foundation. The computations were partially performed at the Poznan Supercomputing and Networking Center (grant number pl0074-02).

## Contributions

P.R. and B.E.D. conceived and designed the study. A.Z. assigned host taxonomy for *Mushuviridae* phages and contributed to figure design. M.S., M.A.A.F. and S.A. conducted transcriptomic and proteomic analyses. P.R. collected and analysed data, designed figures, and wrote the manuscript with substantial contributions from A.Z. and B.E.D. All authors reviewed and approved the final manuscript.

## Extended Data Figures

**Extended Data Fig. 1.**
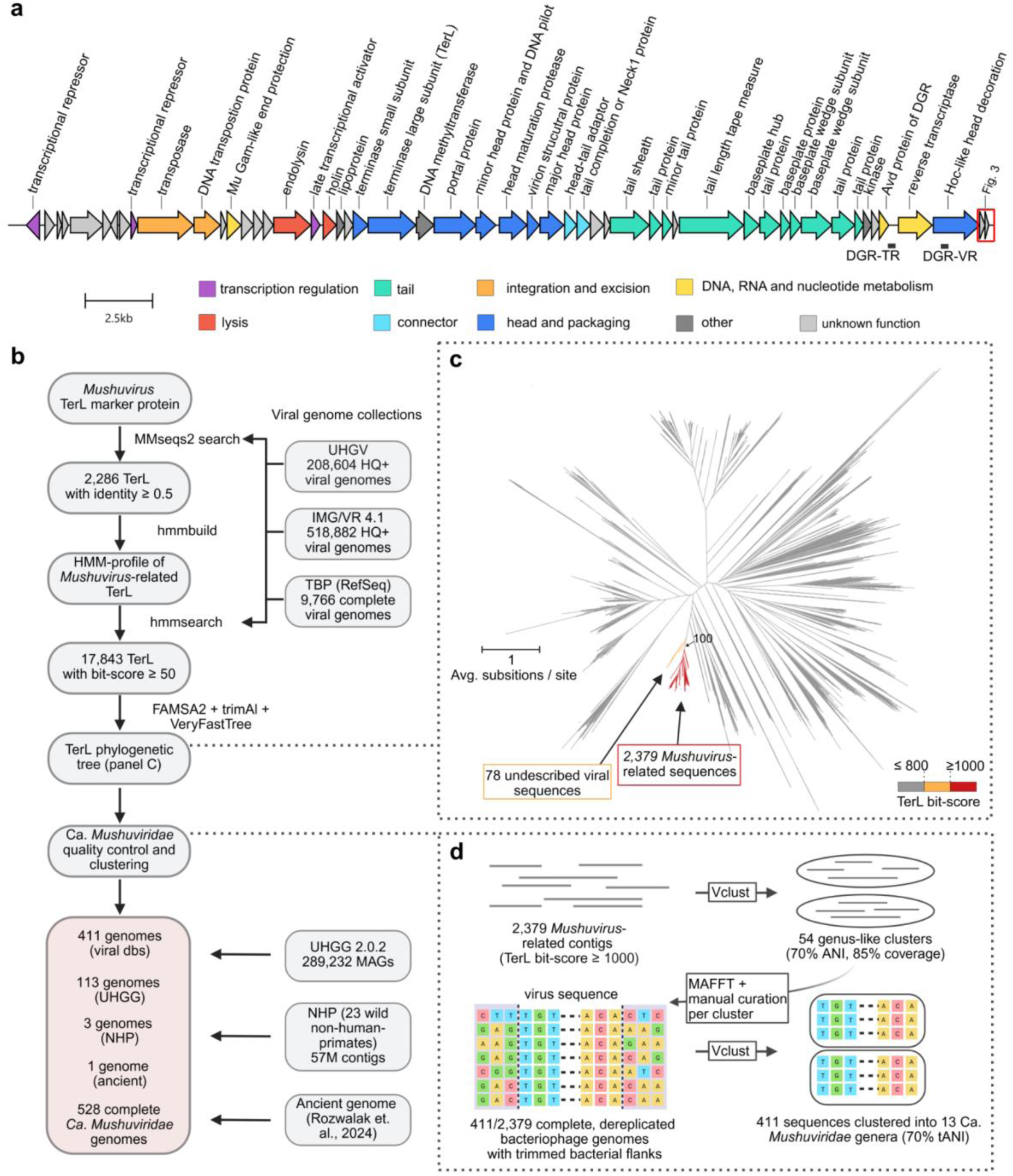
***Mushuviridae* represents a distinct lineage within the global virome a,** Genome architecture of the reference *Mushuvirus mushu* (RefSeq accession: NC_047913.1) functionally annotated using Pharokka^100^ v1.7.314 and Phold^77^ v02.0. Black horizontal lines indicate template (TR) and variable regions (VR) associated with the DGR mechanism, identified using myDGR^99^. The red rectangle highlights two hypothetical proteins presented in detail in Figure 3. **b,** Workflow for the discovery of homologous sequences to the hallmark *Mushuvirus mushu* terminase large subunit (TerL) gene (YP_009797323.1). HQ+: high-quality and complete genomes. **c,** An unrooted approximate maximum-likelihood tree of *Mushuvirus*-related TerL genes and their relatives with bitscore ≥ 50 from hmmsearch against public viral sequence databases. The tree was constructed in VeryFastTree^68^ v3.1.0. The monophyletic lineage of *Mushuvirus*-related-TerL is defined by 1,000 bit-score threshold from hmmsearch. The closest viral lineage to mushuviruses is represented by 78 sequences representing currently undescribed *Enterocloster* viruses. **d,** *Mushuviridae* genome quality-control based on identified conserved genome-end motifs separating complete phage genomes from variable bacterial flanking regions. Complete sequences were clustered to different phage genera using Vclust^60^.

**Extended Data Fig. 2.**
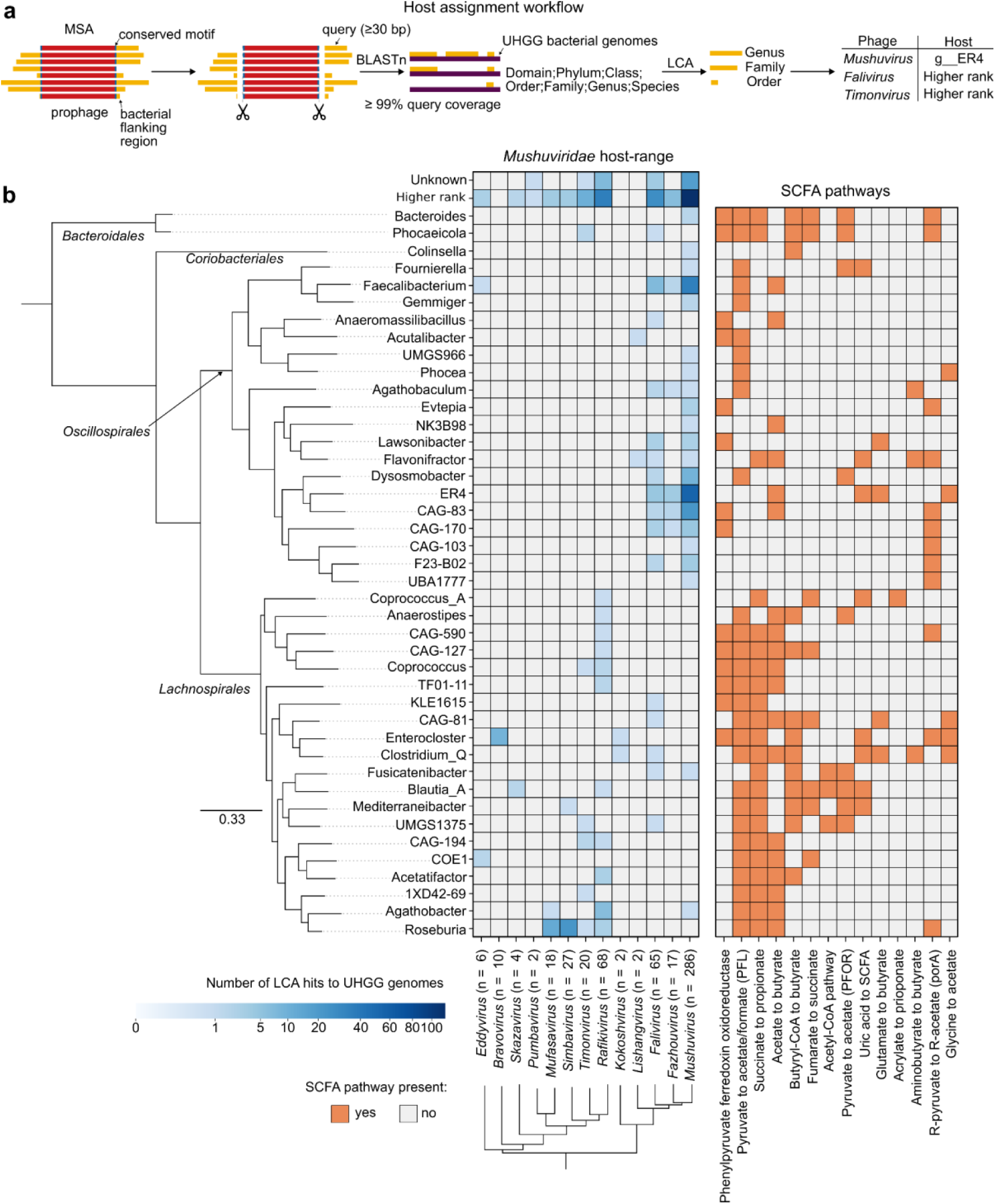
Broad host range of *Mushuviridae* across SCFA-producing bacteria. **(a)** Workflow for host assignment based on the extraction of prophage-flanking bacterial regions. Prophages binned into MAGs or integrated in bacterial isolate chromosomes were assigned to the host without mapping flanking regions. See the Methods section for a detailed description. **(b)** The blue heatmap shows the detected bacteria at the genus rank. The bacterial maximum-likelihood phylogenetic tree was reconstructed from representative UHGG genomes using marker genes (GTDB-Tk^101^). The phage tree at the bottom is a pruned version of the proteomic tree shown in Fig. 2a. The orange heatmap indicates the presence of short-chain fatty acid (SCFA) metabolic pathways detected in representative UHGG genomes (GutSMASH^102^).

**Extended Data Fig. 3.**
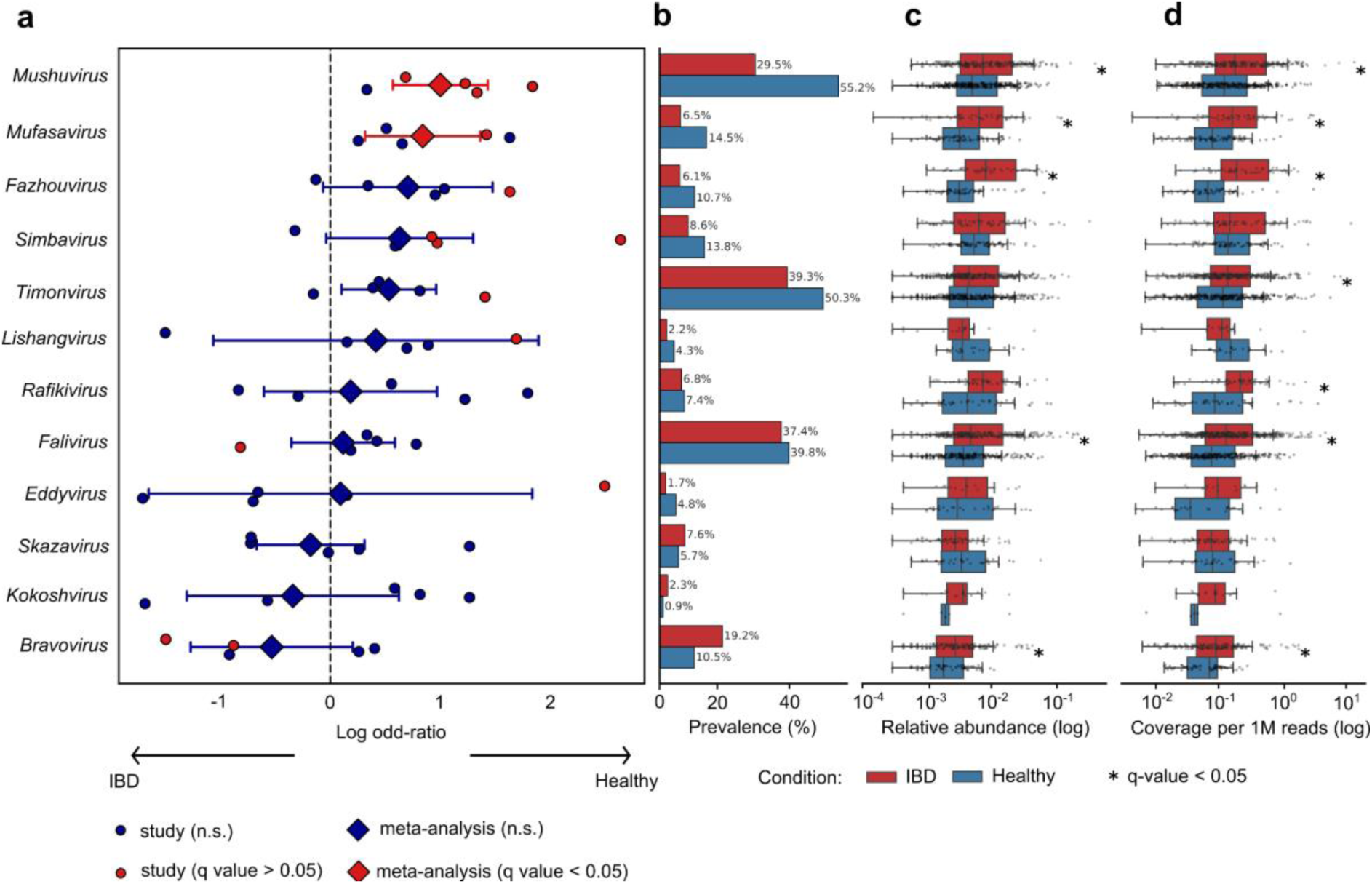
***Mushuviridae* associations with IBD (A)** Forest plot showing the associations of *Mushuviridae* with healthy and inflammatory bowel disease (IBD) patients across 1,448 samples from five studies^84–88^ spanning the USA, China, Denmark, and Spain. Odds ratios were calculated using a pseudocount of 0.5 to prevent division by zero, shown with 95% confidence intervals. Small filled circles represent per-study log odds ratios; large diamonds represent the DerSimonian–Laird random-effects pooled estimate across all studies, with horizontal lines indicating 95% confidence intervals derived from the pooled standard error. Red indicates statistical significance (FDR <5% after Benjamini–Hochberg correction), dark blue indicates non-significance. Positive log OR values indicate depletion in IBD (enrichment in healthy controls); negative values indicate enrichment in IBD.

**Extended Data Fig. 4.**
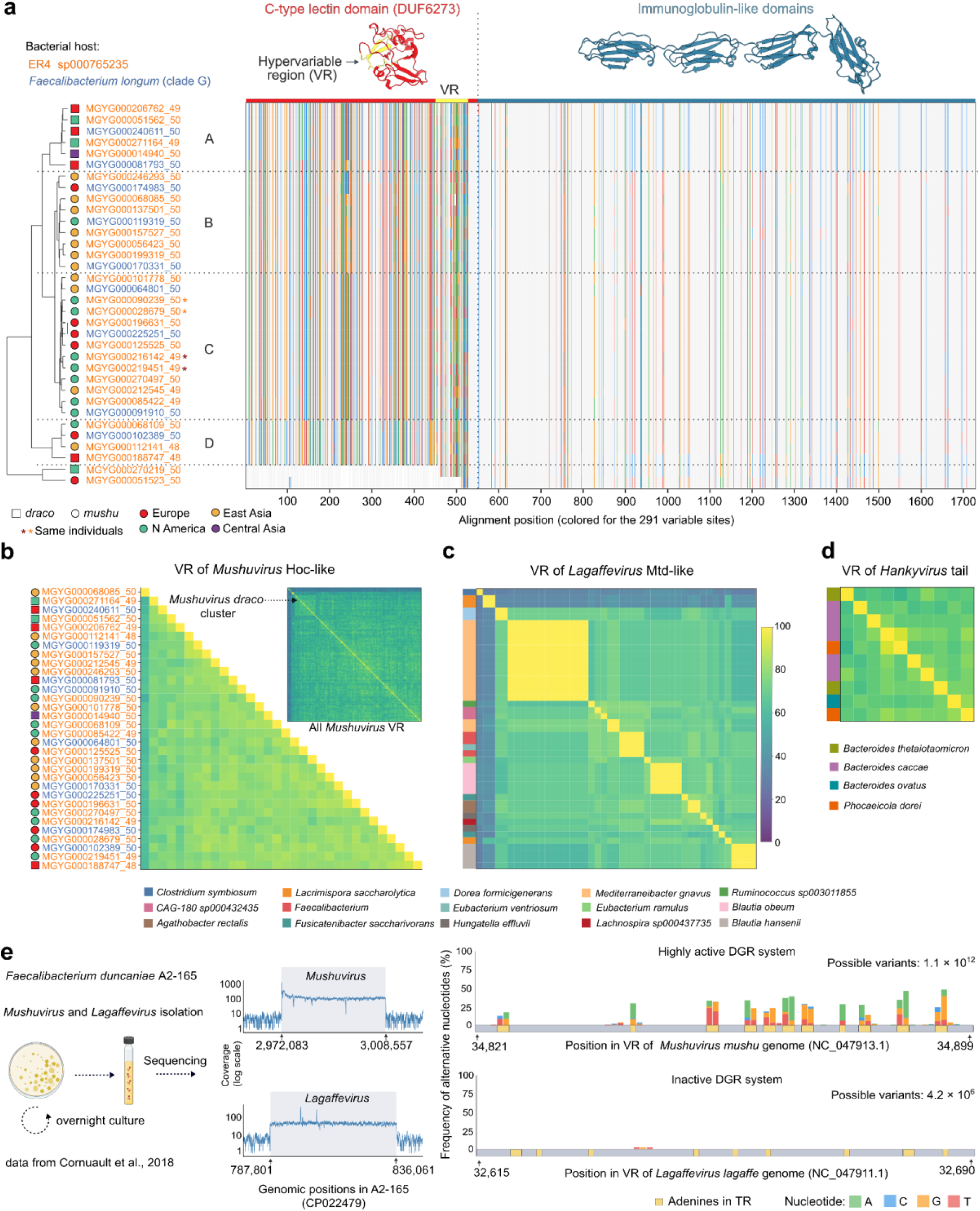
***Mushuvirus* Hoc-like proteins are diversified by a highly active DGR system. a,** Tree based on hierarchical clustering of selected Hoc-like genes from *Mushuvirus* genomes integrated in bacterial MAGs representing ER4 sp000765235 and *Faecalibacterium longum* (clade G). The tree is annotated by taxonomy (marker shape), host (text colour), and geographic location (marker colour) together with multiple sequence alignment (MAFFT^70^) of respective genes. Colored lines indicate positions where the nucleotide in at least one sequence differs from the others. Protein structures were predicted and visualised (AphaFold3) into two, artificially split domains. Asterisks indicate two pairs of sequences coming from the same individuals collected in longitudinal studies. **b,** Global pairwise alignment comparison of *Mushuvirus* amino acid sequences from the short DGR-mutagenised region (VR). **c,** Global pairwise alignment comparison of the VR located in the *Lagaffevirus lagaffe* protein, which is similar to the major tropism determinant (Mtd) encoded by Bordetella phage BPP-1 and plays a role in host switching. *Lagaffevirus* sequences were identified and extracted from multiple UHGG MAGs. **d,** Global pairwise alignment comparison of the VR located in a tail protein of *Hankyvirus* phages identified in *Bacteroides* and *Phocaeicola* bacterial isolates^19^. **e,** After overnight cultivation of *Faecalibacterium duncaniae* strain A2-165, phage particles were isolated from the supernatant and sequenced, revealing that *Mushuvirus mushu* and *Lagaffevirus lagaffe* are active viruses and that *Mushuvirus* possesses a highly active DGR system. DGR activity was measured based on a reanalysis of data from experiments published by Cornuault et al., 2018^22^.

