## Supplementary Figures for "Cosmopolitan gut bacteriophages expand the phenotype of health-related bacteria"

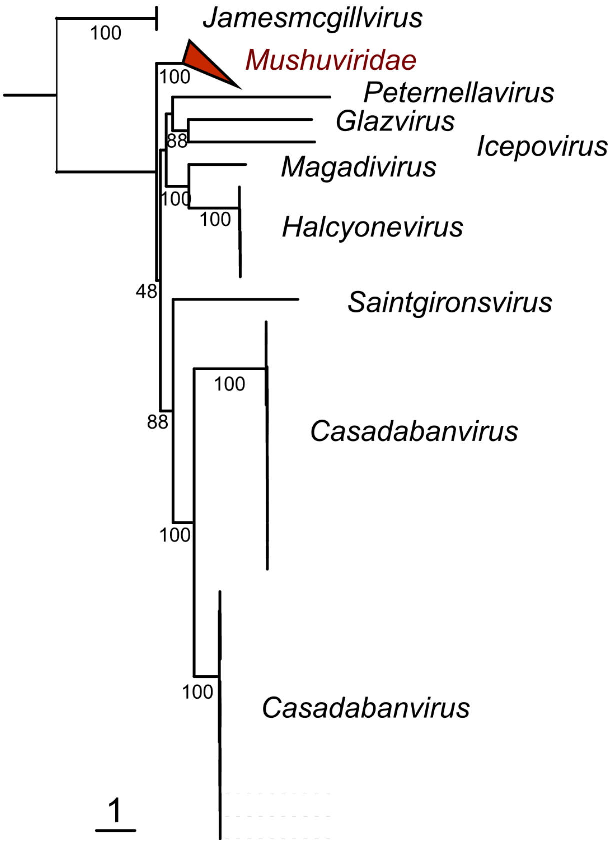


**Supplementary Figure 1. Maximum likelihood phylogenetic tree of *Mushuviridae* and the most closely related bacteriophages based on the Terminase Large Subunit (TerL).** Sequences used to build the tree were selected from 6,741 International Committee on Taxonomy of Viruses (ICTV)-classified bacteriophages (Virus Metadata Resource, VMR v40.1) using an HMM profile specific for *Mushuvirus* TerL-related sequences (Methods; Identification of *Mushuviridae* sequences). Selected TerL proteins were aligned using FAMSA[1] v2.2.3, and the tree was constructed with IQ-TREE v2.1.4. The tree is midpoint-rooted, and node labels indicate bootstrap support.

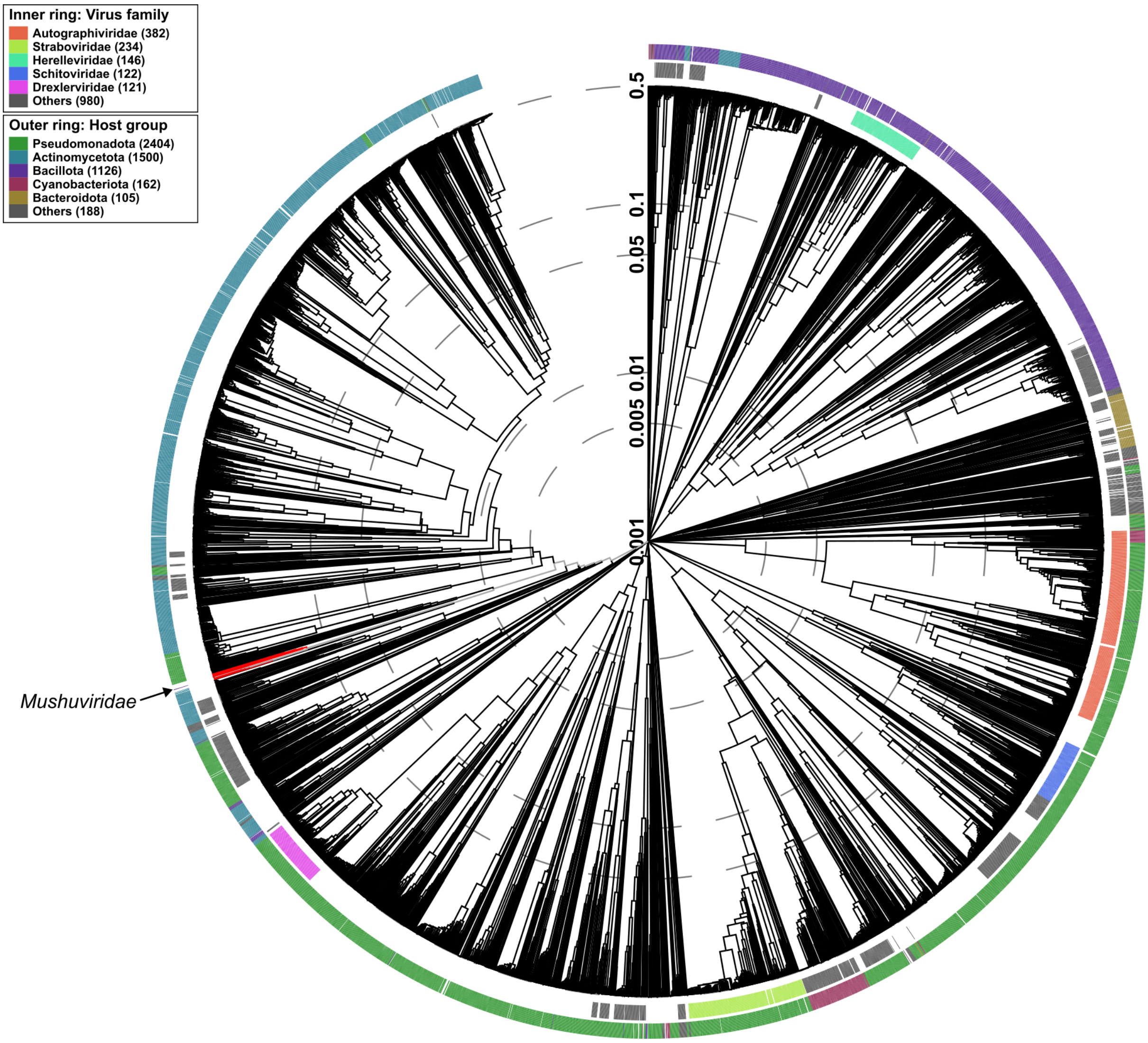


**Supplementary Figure 2. Proteome-based phylogenetic placement of 13 representative *Mushuviridae* genomes in the context of 5,632 bacteriophages from RefSeq v220.** The tree was generated using the VipTree webservice (https://www.genome.jp/viptree/). The *Mushuviridae* clade (highlighted in red and indicated by an arrow) is phylogenetically closer to phages infecting *Pseudomonadota* and *Actinomycetota* than to other *Bacillota*-infecting phages. The inner ring denotes viral family assignments, while the outer ring represents host phylum associations.

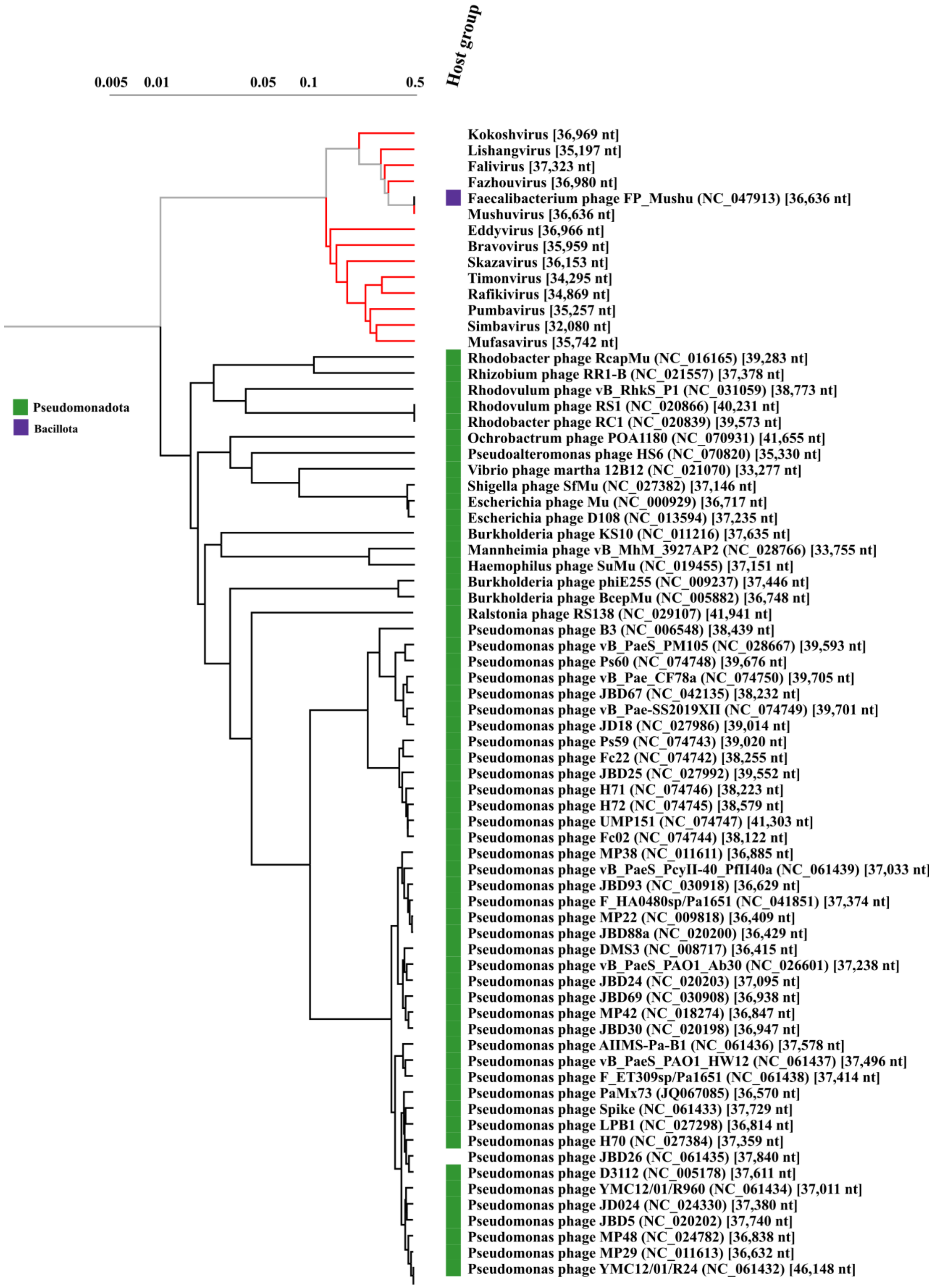


**Supplementary Figure 3. Phylogenetic relationship between *Mushuviridae* and their closest relatives.** This subset of the global VipTree highlights the sister-clade relationship between *Mushuviridae* (red branches) and various Mu-like phages (black branches). Host phylum associations are indicated by the colour bar, showing *Mushuviridae* primarily associated with *Bacillota* (purple) while the neighbouring Mu-like phages are associated with *Pseudomonadota* (green). Scale bar represents genomic distance based on proteome-wide similarity.

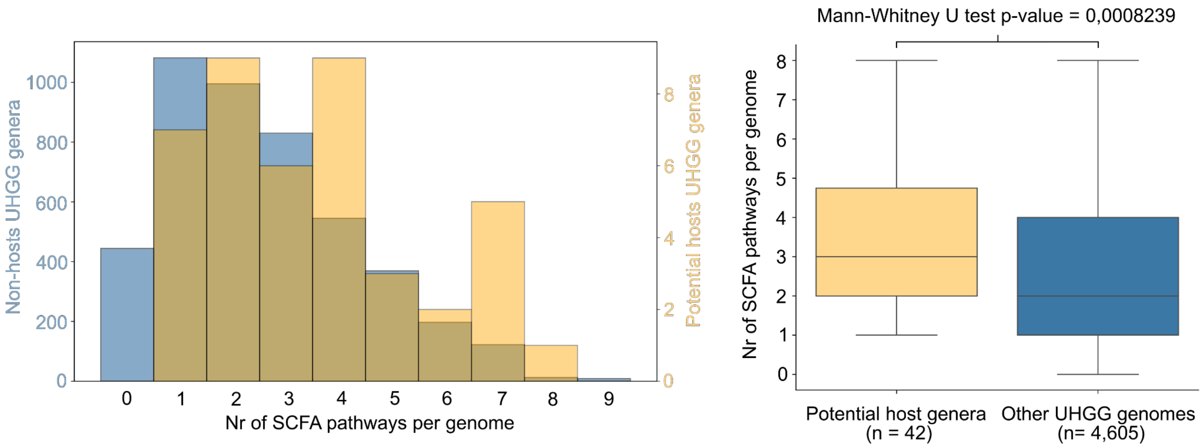


**Supplementary Figure 4. Enrichment of short-chain fatty acid (SCFA) metabolic pathways in the predicted hosts of *Mushuviridae*.** SCFA pathways were annotated using gutSMASH[2] in representative genomes from the Unified Human Gastrointestinal Genome (UHGG) database v2.0.2. The histogram (left) and boxplot (right) compare the number of metabolic pathways per genome between identified potential host genera ($n$ = 42) and non-host UHGG genomes ($n$ = 4,605). Statistical significance was determined using a two-sided Mann-Whitney U test ($p$ = 0.0008239), indicating that *Mushuviridae* hosts carry a significantly higher number of SCFA-producing pathways compared to other human gut bacteria.

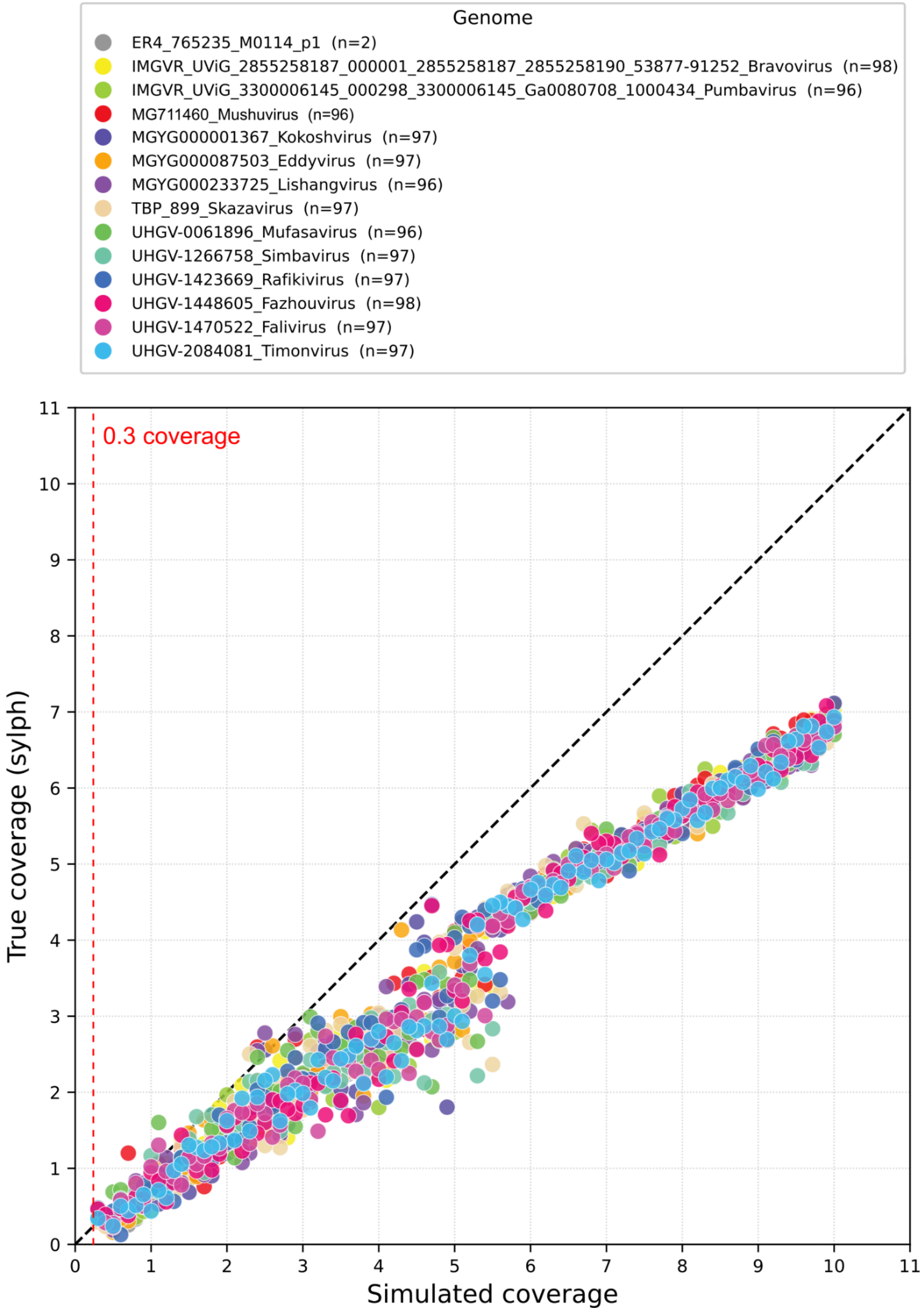


**Supplementary Figure 5. Validation of sylph[4] taxonomic profiling using 100 simulated metagenomes generated in InSilicoSeq[3].** The plot compares known (simulated) *Mushuviridae* coverage against the coverage estimated by sylph ("True coverage"). The dashed red line indicates the 0.3× detection threshold. Sylph showed high precision for low-abundant viral genomes, with only two instances of missclassification of *Mushuviridae* as ER4. The observed decline in accuracy at coverage levels above 3× likely results from a switch in sylph's internal coverage estimator, which depends on the median *k*-mer coverage.

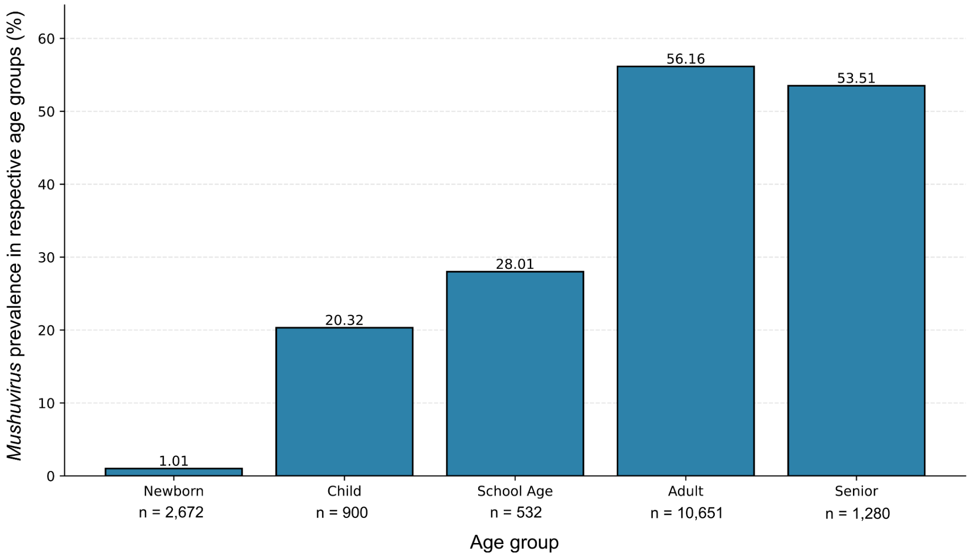


**Supplementary Figure 6. Prevalence of *Mushuvirus* across human life stages.** Taxonomic profiles were generated using sylph[4] and integrated with age metadata from CuratedMetagenomicData[5] v3.21. The bar chart illustrates a successive increase in *Mushuvirus* prevalence starting from newborns (1.01%) through childhood and school age, peaking in adults (56.16%). Sample sizes (*n*) for each age group are provided below the x-axis labels.

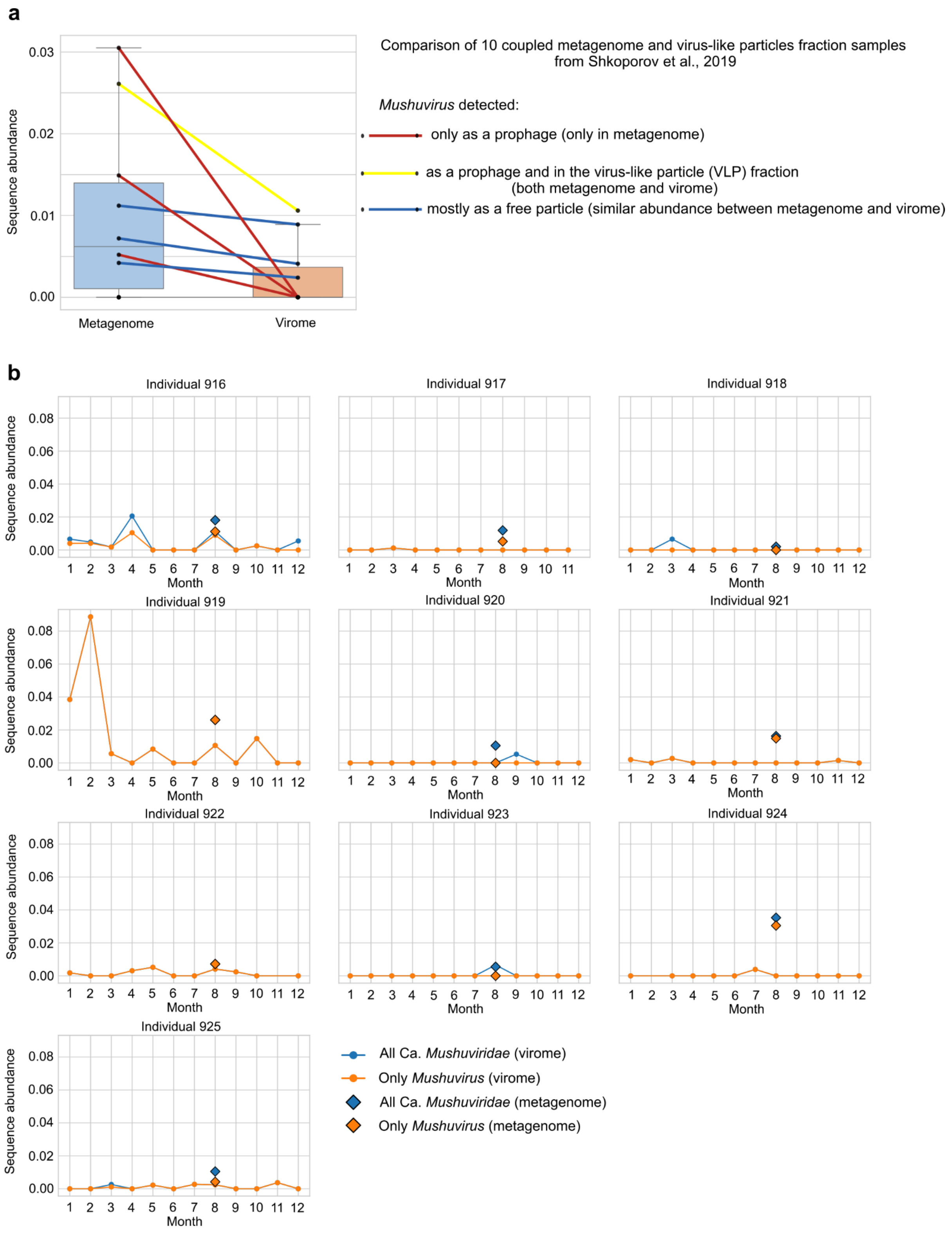


**Supplementary Figure 7. Evidence of active prophage induction within the *Mushuviridae*.** Sequence abundance for both panels was calculated with sylph[4] based on coupled metagenome and virus-like particle (VLP) fraction samples from Shkoporov et al., 2019[6]. **a,** Comparison of 10 coupled samples demonstrating that *Mushuvirus* occurs as an integrated prophage (detected only in the metagenome), as an active particle undergoing induction (detected in both fractions), or predominantly as a free particle. **b,** Time-series analysis of sequence abundance for all *Mushuviridae* and M*ushuvirus* phages across 10 individuals over 12 months (see Taxonomic profiling in Methods). Diamonds represent metagenomic abundance at month 8, while lines represent longitudinal virome abundance. Values are presented as sequence abundance to allow for direct comparison across identical genomes.

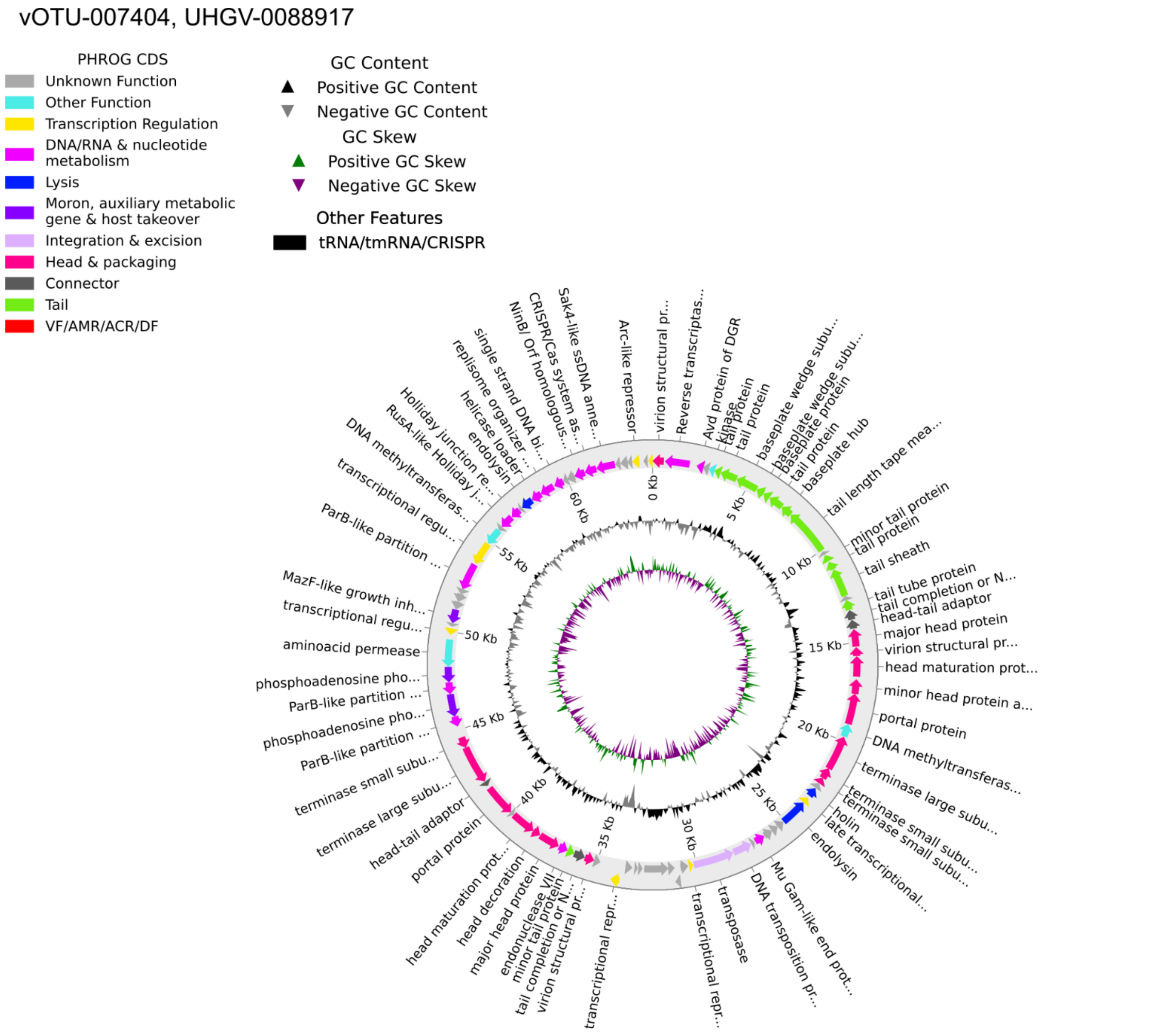


| | Description | Scientific Name | Max Score | Total Score | Query Cover | E value | Per. Ident | Acc. Len | Accession |
|---|---|---|---|---|---|---|---|---|---|
| ☑ | Faecalibacterium duncaniae strain JCM 31915 chromosome, complete genome | Faecalibacteriu... | 56567 | 57867 | 53% | 0.0 | 96.93% | 3102523 | CP048437.1 |
| ☑ | Faecalibacterium phage FP_Mushu, complete genome | Faecalibacteriu... | 56567 | 57865 | 53% | 0.0 | 96.93% | 36636 | NC_047913.1 |
| ☑ | Faecalibacterium duncaniae strain A2165 chromosome, complete genome | Faecalibacteriu... | 56560 | 57859 | 54% | 0.0 | 96.93% | 3110044 | CP022479.1 |
| ☑ | TPA_asm: Mushuvirus mushu strain Zape3, complete genome | Mushuvirus mushu | 55081 | 56359 | 53% | 0.0 | 96.15% | 36623 | BK063464.1 |
| ☑ | Faecalibacterium sp. i25-0019-C1 DNA, complete genome | Faecalibacteriu... | 41790 | 55438 | 52% | 0.0 | 96.30% | 2876947 | AP031428.1 |
| ☑ | MAG TPA_asm: Caudoviricetes sp. isolate ctDPK14, partial genome | Caudoviricetes sp. | 21195 | 31193 | 44% | 0.0 | 86.48% | 56390 | BK046128.1 |
| ☑ | MAG TPA_asm: Caudoviricetes sp. isolate ctL9011, partial genome | Caudoviricetes sp. | 20192 | 20192 | 19% | 0.0 | 96.06% | 39429 | BK021889.1 |
| ☑ | MAG TPA_asm: Caudoviricetes sp. isolate ctLsJ4, partial genome | Caudoviricetes sp. | 20138 | 20138 | 18% | 0.0 | 97.61% | 21050 | BK016891.1 |
| ☑ | MAG TPA_asm: Caudoviricetes sp. isolate ctnte1, partial genome | Caudoviricetes sp. | 17544 | 45075 | 45% | 0.0 | 95.13% | 53749 | BK052196.1 |
| ☑ | MAG TPA_asm: Caudoviricetes sp. isolate ctDG41, partial genome | Caudoviricetes sp. | 15538 | 47457 | 45% | 0.0 | 97.56% | 55070 | BK038158.1 |
| ☑ | MAG TPA_asm: Caudoviricetes sp. isolate ctfpA15, partial genome | Caudoviricetes sp. | 14885 | 17744 | 22% | 0.0 | 95.24% | 31145 | BK032468.1 |
| ☑ | Unidentified phage clone 2209_scaffold1451 genomic sequence | Bacteriophage sp. | 11960 | 30774 | 26% | 0.0 | 99.82% | 19156 | JQ680376.1 |
| ☑ | MAG TPA_asm: Caudoviricetes sp. isolate ctFby16, partial genome | Caudoviricetes sp. | 10988 | 19626 | 43% | 0.0 | 78.80% | 36759 | BK033876.1 |
| ☑ | MAG: Caudoviricetes sp. isolate MSP1624 genomic sequence | Caudoviricetes sp. | 10922 | 16627 | 42% | 0.0 | 77.36% | 43002 | OR221460.1 |
| ☑ | MAG TPA_asm: Bacteriophage sp. isolate ctvpl5, partial genome | Bacteriophage sp. | 10599 | 19361 | 43% | 0.0 | 78.37% | 44941 | BK055849.1 |
| ☑ | MAG TPA_asm: Caudoviricetes sp. isolate ctdux2, partial genome | Caudoviricetes sp. | 8486 | 19450 | 21% | 0.0 | 90.60% | 14142 | BK021526.1 |
| ☑ | Unidentified phage clone 1013_scaffold7838 genomic sequence | Bacteriophage sp. | 6721 | 6721 | 7% | 0.0 | 92.39% | 6416 | JQ680355.1 |
| ☑ | MAG TPA_asm: Caudovirales sp. ctNZz8, partial genome | Caudovirales sp.... | 6656 | 21594 | 34% | 0.0 | 90.64% | 53858 | BK015770.1 |
| ☑ | MAG TPA_asm: Caudoviricetes sp. isolate ctXKr2, partial genome | Caudoviricetes sp. | 6510 | 6510 | 16% | 0.0 | 78.43% | 14576 | BK042658.1 |
| ☑ | MAG TPA_asm: Caudoviricetes sp. isolate ctGyO1, partial genome | Caudoviricetes sp. | 6242 | 6242 | 8% | 0.0 | 89.63% | 25703 | BK032034.1 |
| ☑ | MAG TPA_asm: Caudoviricetes sp. isolate ctTMU2, partial genome | Caudoviricetes sp. | 4710 | 13314 | 30% | 0.0 | 81.74% | 36388 | BK025231.1 |
| ☑ | Agathobaculum sp. NTUH-O15-33 chromosome, complete genome | Agathobaculum ... | 3587 | 12969 | 38% | 0.0 | 78.52% | 4005854 | CP136187.1 |
| ☑ | MAG TPA_asm: Caudoviricetes sp. isolate ctRdn3, partial genome | Caudoviricetes sp. | 3533 | 9171 | 25% | 0.0 | 76.47% | 16816 | BK053996.1 |

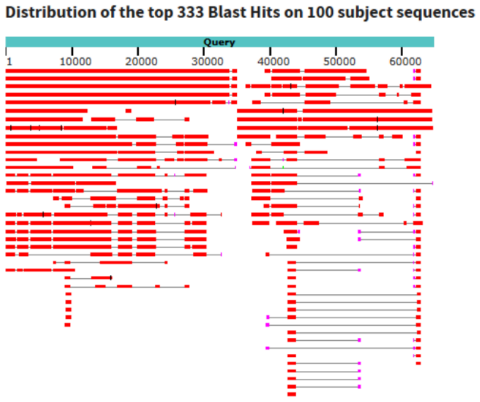


**Supplementary Figure 8. Characterisation of the chimeric representative genome UHGV-0088917 (vOTU-007404) from the UHGV catalogue**[7]. The circular map (top) illustrates the 64 kb sequence, which is a phage–phage chimera with the core *Mushuvirus* modules alongside an additional ~28 kb phage-derived region. Functional annotation of PHROG CDS reveals distinct structural, replication, and lysis modules, as well as GC content and skew profiles. Despite its chimeric nature, the sequence was classified as a complete genome by CheckV[8] due to the presence of viral hallmark genes throughout. The BLAST results and hit distribution (bottom) confirm the high sequence identity with *Faecalibacterium* phages and the segmented nature of the assembly relative to other *Caudoviricetes* isolates.

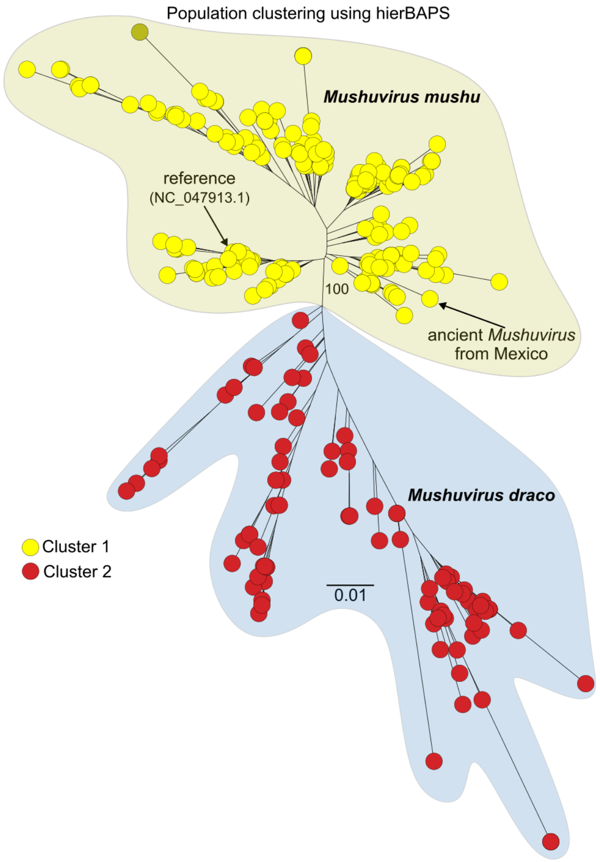


**Supplementary Figure 9. Species delineation of a global *Mushuvirus* population via Bayesian model-based clustering.** Hierarchical clustering of DNA sequence data was performed using the RhierBAPS R package[9] to identify nested genetic population structures within the *Mushuvirus* collection. The analysis supports the separation of the global population into two distinct clusters corresponding to *Mushuvirus mushu* (Cluster 1, yellow) and *Mushuvirus draco* (Cluster 2, red). Key sequences are highlighted, including the reference genome (NC_047913.1) and an ancient *Mushuvirus* sequence from Mexico (BK063464). The scale bar represents substitutions per site, and the central node indicates bootstrap support.

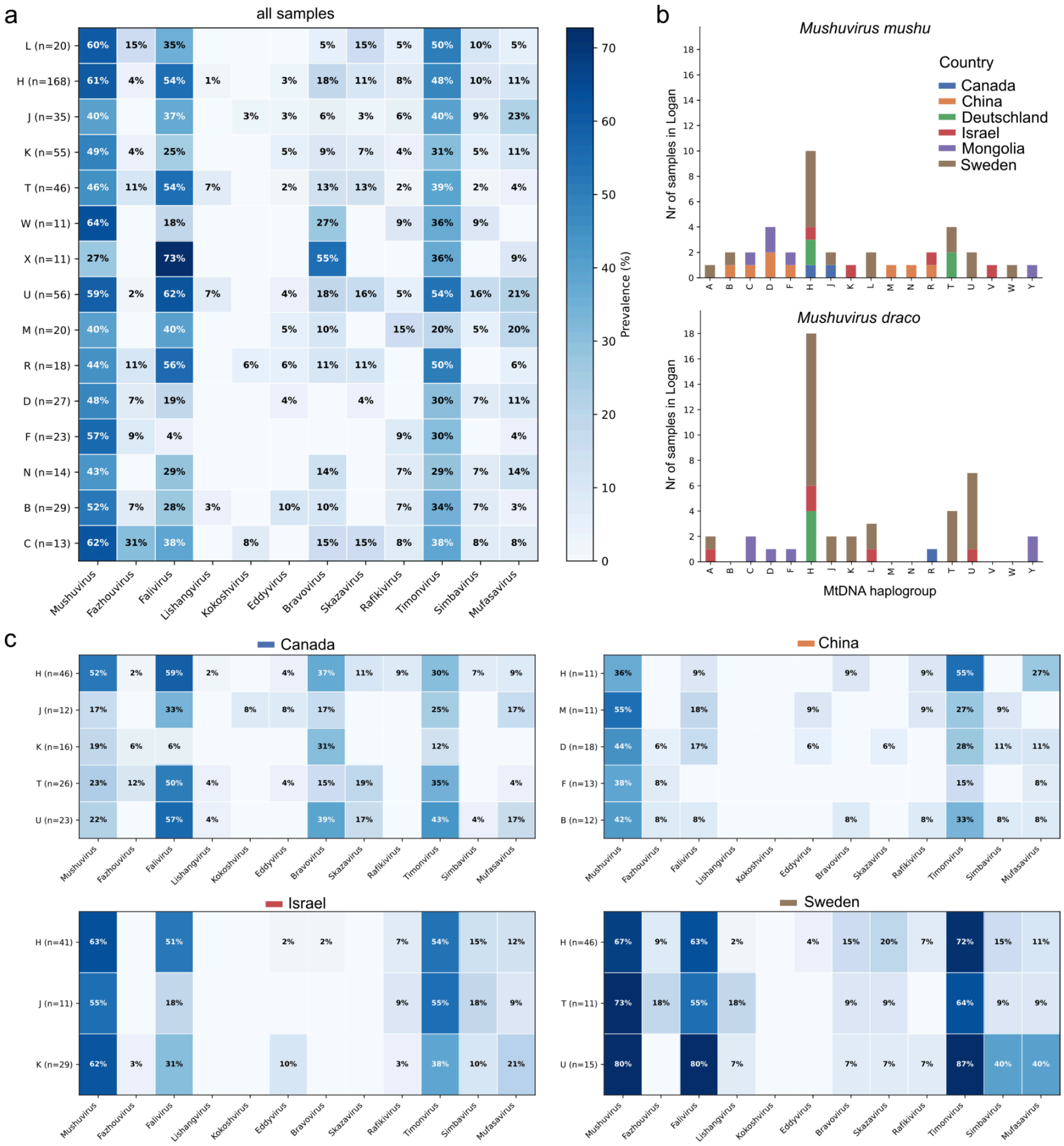


**Supplementary Figure 10. Prevalence of *Mushuviridae* across human mitochondrial DNA (mtDNA) haplogroups.** mtDNA haplogroups were inferred from stool metagenome sequences (described in Sarhan et al., 2025[11]) and linked to *Mushuviridae* occurrence based on shared sample identifiers. **a,** Heatmap showing the prevalence of various *Mushuviridae* genera across 508 samples with successful haplogroup assignment. **b,** Global distribution and occurrence of *Mushuvirus mushu* and *Mushuvirus draco* across different haplogroups and countries, determined using diagnostic sequence searches in the Logan database. **c,** Country-specific heatmaps demonstrating consistent enrichment of *Falivirus* in the West Eurasian H haplogroup and relative depletion in the East Asian F haplogroup across Canada, China, Israel, and Sweden.

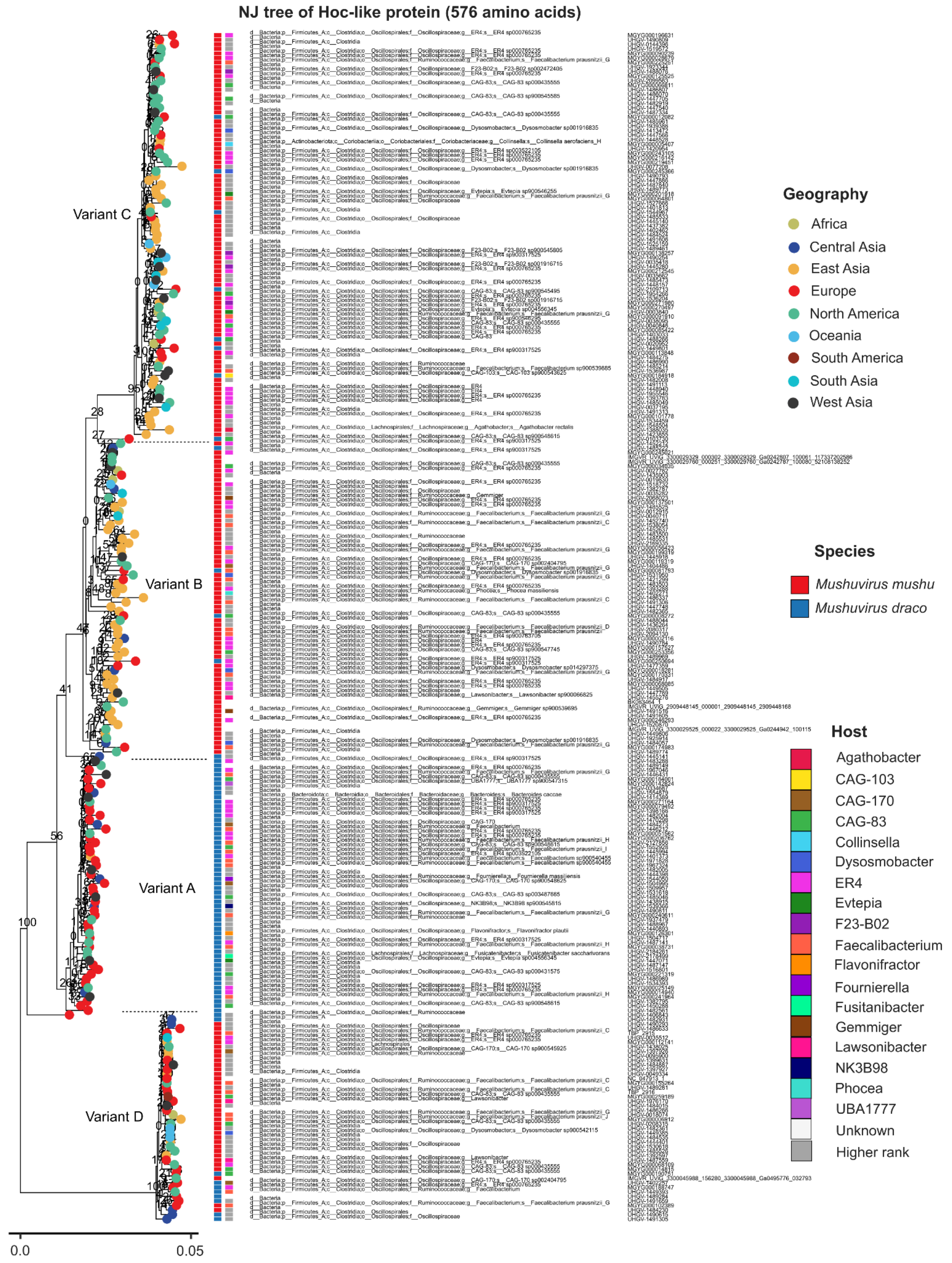


**Supplementary Figure 11. Neighbour-joining phylogenetic tree of Hoc-like proteins revealing four geographically structured variants (Variants A–D).** Multiple sequence alignment and tree inference were performed using the MAFFT web service with default parameters and 100 bootstrap replicates. The tree illustrates the genetic diversity of Hoc-like proteins across *Mushuvirus mushu* and *Mushuvirus draco*, host genera, and global geographic regions. Nodes are colour-coded by geography, and vertical bars indicate species and host associations. The scale bar represents amino acid substitutions per site.

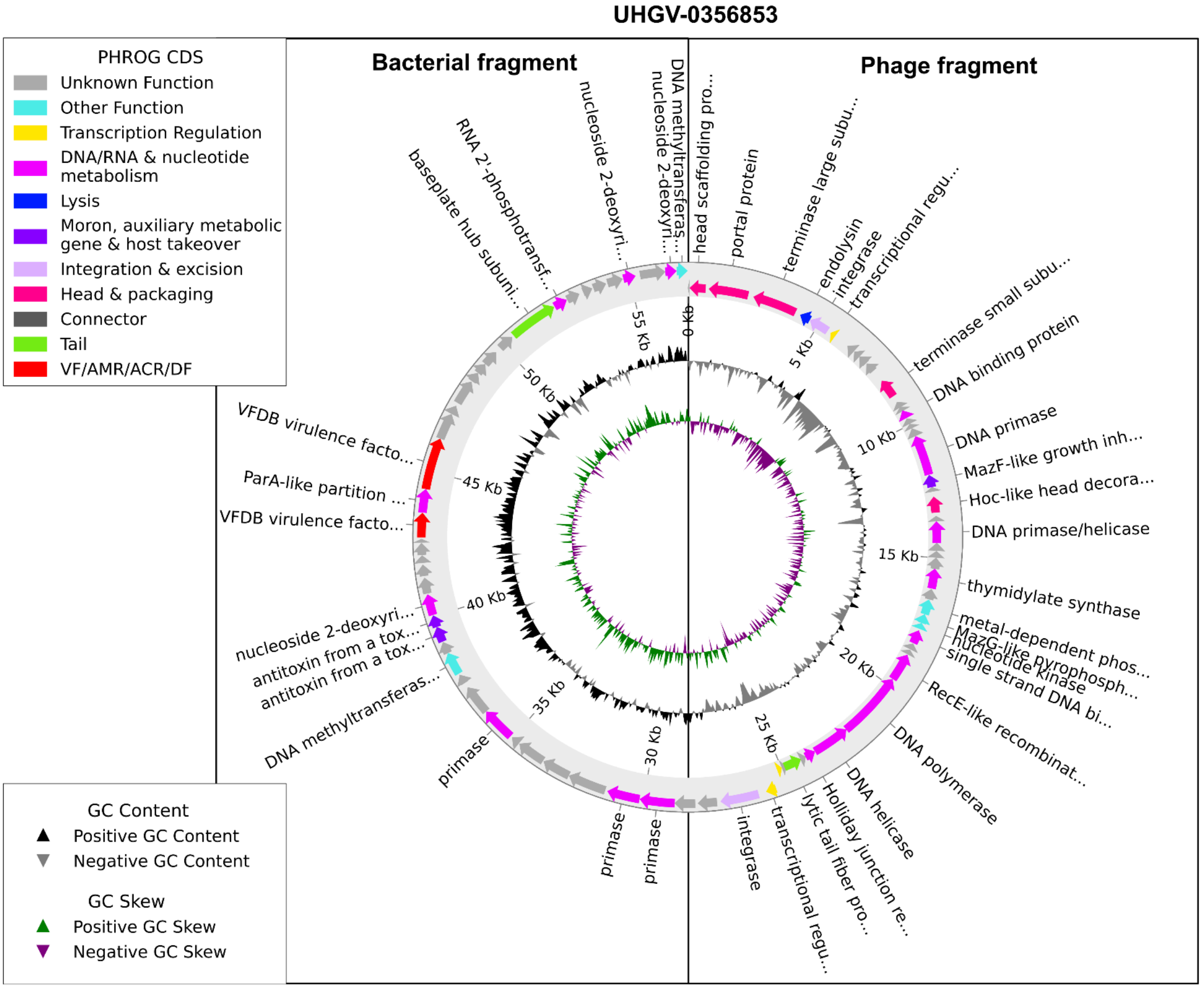


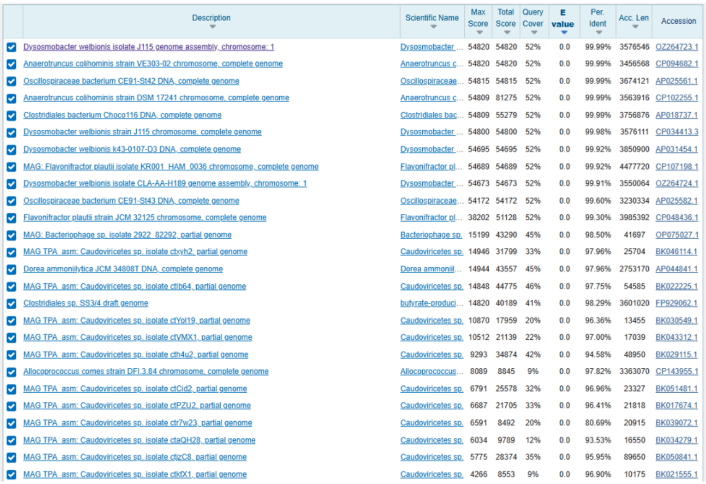

| Description | Scientific Name | Max Score | Total Score | Query Cover | E value | Per. Ident | Acc. Len | Accession |
|---|---|---|---|---|---|---|---|---|
| Dysosmobacter welbionis isolate J115 genome assembly, chromosome: 1 | Dysosmobacter ... | 54820 | 54820 | 52% | 0.0 | 99.99% | 3576546 | OZ264723.1 |
| Anaerotruncus colihominis strain VE303-02 chromosome, complete genome | Anaerotruncus c... | 54820 | 54820 | 52% | 0.0 | 99.99% | 3456568 | CP094682.1 |
| Oscillospiraceae bacterium CE91-St42 DNA, complete genome | Oscillospiraceae... | 54815 | 54815 | 52% | 0.0 | 99.99% | 3674121 | AP025561.1 |
| Anaerotruncus colihominis strain DSM 17241 chromosome, complete genome | Anaerotruncus c... | 54809 | 81275 | 52% | 0.0 | 99.99% | 3563916 | CP102255.1 |
| Clostridiales bacterium Choco116 DNA, complete genome | Clostridiales bac... | 54809 | 55279 | 52% | 0.0 | 99.99% | 3756876 | AP018737.1 |
| Dysosmobacter welbionis strain J115 chromosome, complete genome | Dysosmobacter ... | 54800 | 54800 | 52% | 0.0 | 99.98% | 3576111 | CP034413.3 |
| Dysosmobacter welbionis k43-0107-D3 DNA, complete genome | Dysosmobacter ... | 54695 | 54695 | 52% | 0.0 | 99.92% | 3850900 | AP031454.1 |
| MAG: Flavonifractor plautii isolate KR001_HAM_0036 chromosome, complete genome | Flavonifractor pl... | 54689 | 54689 | 52% | 0.0 | 99.92% | 4477720 | CP107198.1 |
| Dysosmobacter welbionis isolate CLA-AA-H189 genome assembly, chromosome: 1 | Dysosmobacter ... | 54673 | 54673 | 52% | 0.0 | 99.91% | 3550064 | OZ264724.1 |
| Oscillospiraceae bacterium CE91-St43 DNA, complete genome | Oscillospiraceae... | 54172 | 54172 | 52% | 0.0 | 99.60% | 3230334 | AP025582.1 |
| Flavonifractor plautii strain JCM 32125 chromosome, complete genome | Flavonifractor pl... | 38202 | 51128 | 52% | 0.0 | 99.30% | 3985392 | CP048436.1 |
| MAG: Bacteriophage sp. isolate 2922_82292, partial genome | Bacteriophage sp. | 15199 | 43290 | 45% | 0.0 | 98.50% | 41697 | OP075027.1 |
| MAG TPA_asm: Caudoviricetes sp. isolate ctxyh2, partial genome | Caudoviricetes sp. | 14946 | 31799 | 33% | 0.0 | 97.96% | 25704 | BK046114.1 |
| Dorea ammoniilytica JCM 34808T DNA, complete genome | Dorea ammoniil... | 14944 | 43557 | 45% | 0.0 | 97.96% | 2753170 | AP044841.1 |
| MAG TPA_asm: Caudoviricetes sp. isolate ctIb64, partial genome | Caudoviricetes sp. | 14848 | 44775 | 46% | 0.0 | 97.75% | 54585 | BK022225.1 |
| Clostridiales sp. SS3/4 draft genome | butyrate-produci... | 14820 | 40189 | 41% | 0.0 | 98.29% | 3601020 | FP929062.1 |
| MAG TPA_asm: Caudoviricetes sp. isolate ctYol19, partial genome | Caudoviricetes sp. | 10870 | 17959 | 20% | 0.0 | 96.36% | 13455 | BK030549.1 |
| MAG TPA_asm: Caudoviricetes sp. isolate ctVMX1, partial genome | Caudoviricetes sp. | 10512 | 21139 | 22% | 0.0 | 97.00% | 17039 | BK043312.1 |
| MAG TPA_asm: Caudoviricetes sp. isolate cth4u2, partial genome | Caudoviricetes sp. | 9293 | 34874 | 42% | 0.0 | 94.58% | 48950 | BK029115.1 |
| Allocoprococcus comes strain DFI.3.84 chromosome, complete genome | Allocoprococcus... | 8089 | 8845 | 9% | 0.0 | 97.82% | 3363070 | CP143955.1 |
| MAG TPA_asm: Caudoviricetes sp. isolate ctCid2, partial genome | Caudoviricetes sp. | 6791 | 25578 | 32% | 0.0 | 96.96% | 23327 | BK051481.1 |
| MAG TPA_asm: Caudoviricetes sp. isolate ctPZU2, partial genome | Caudoviricetes sp. | 6687 | 21705 | 33% | 0.0 | 96.41% | 21818 | BK017674.1 |
| MAG TPA_asm: Caudoviricetes sp. isolate ctr7w23, partial genome | Caudoviricetes sp. | 6591 | 8492 | 20% | 0.0 | 80.69% | 20915 | BK039072.1 |
| MAG TPA_asm: Caudoviricetes sp. isolate ctaQH28, partial genome | Caudoviricetes sp. | 6034 | 9789 | 12% | 0.0 | 93.53% | 16550 | BK034279.1 |
| MAG TPA_asm: Caudoviricetes sp. isolate ctjzC8, partial genome | Caudoviricetes sp. | 5775 | 28374 | 35% | 0.0 | 95.95% | 89650 | BK050841.1 |
| MAG TPA_asm: Caudoviricetes sp. isolate ctkfX1, partial genome | Caudoviricetes sp. | 4266 | 8553 | 9% | 0.0 | 96.90% | 10175 | BK021555.1 |

### Bacterial fragment

**Distribution of the top 21 Blast Hits on 11 subject sequences**

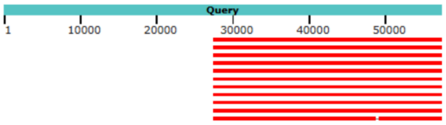


### Phage fragment

**Distribution of the top 90 Blast Hits on 12 subject sequences**

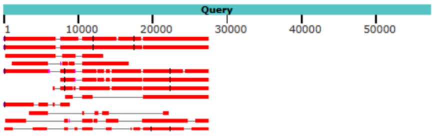

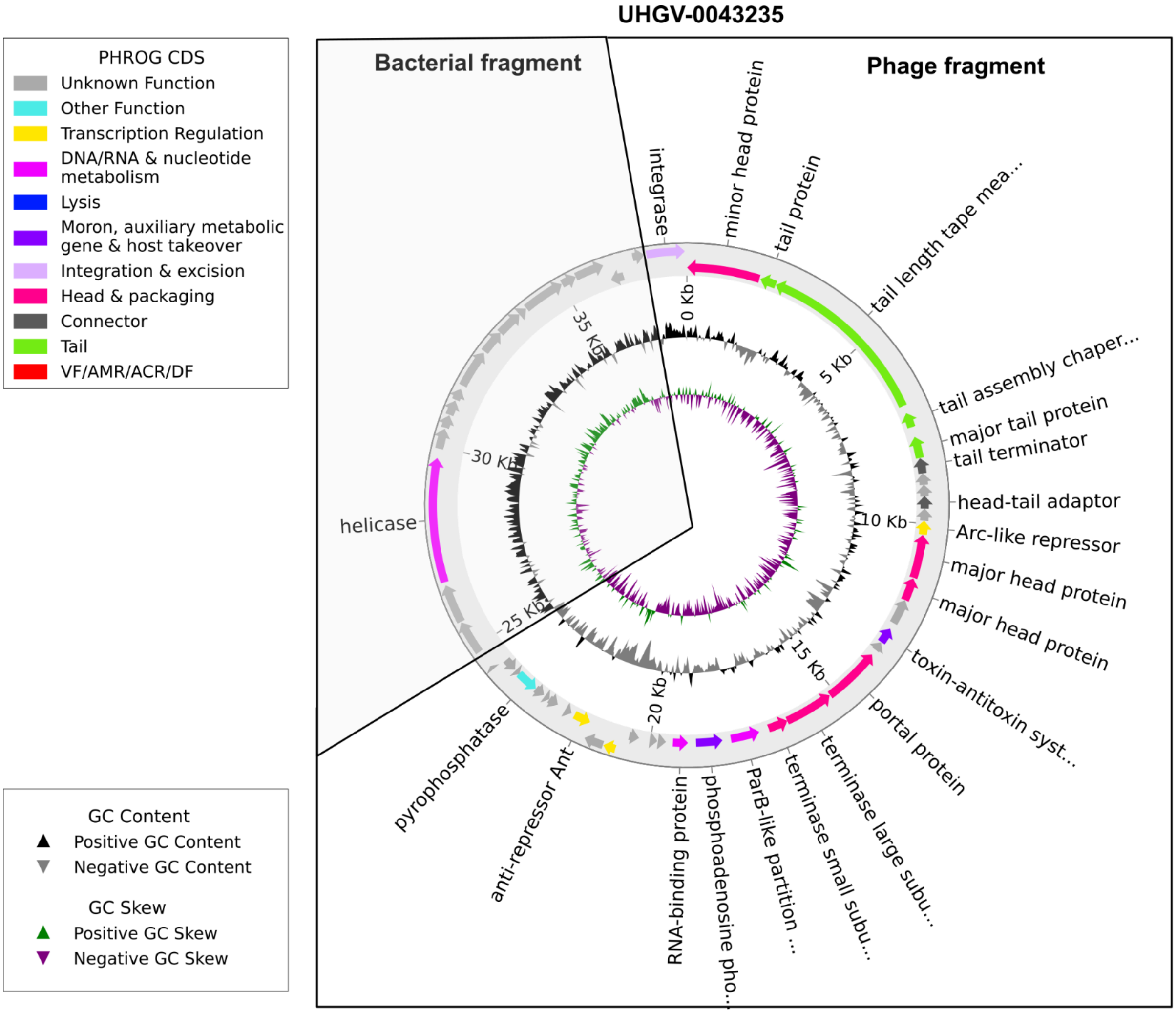


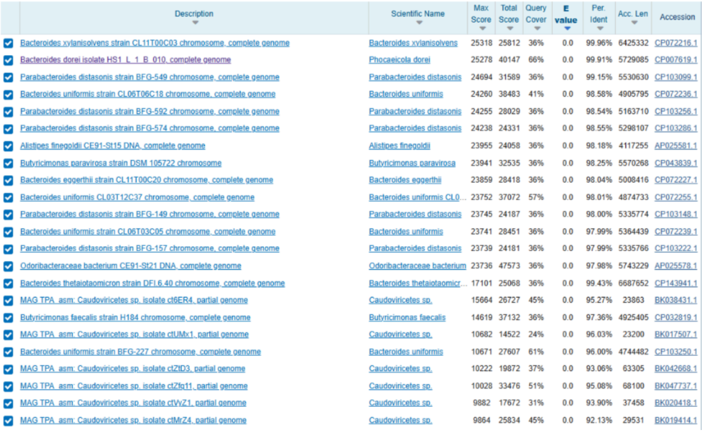

| | Description | Scientific Name | Max Score | Total Score | Query Cover | E value | Per. Ident | Acc. Len | Accession |
|---|---|---|---|---|---|---|---|---|---|
| ☑ | Bacteroides xylanisolvens strain CL11T00C03 chromosome, complete genome | Bacteroides xylanisolvens | 25318 | 25812 | 36% | 0.0 | 99.96% | 6425332 | CP072216.1 |
| ☑ | Bacteroides dorei isolate HS1_L_1_B_010, complete genome | Phocaeicola dorei | 25278 | 40147 | 66% | 0.0 | 99.91% | 5729085 | CP007619.1 |
| ☑ | Parabacteroides distasonis strain BFG-549 chromosome, complete genome | Parabacteroides distasonis | 24694 | 31589 | 36% | 0.0 | 99.15% | 5530630 | CP103099.1 |
| ☑ | Bacteroides uniformis strain CL06T06C18 chromosome, complete genome | Bacteroides uniformis | 24260 | 38483 | 41% | 0.0 | 98.58% | 4905795 | CP072236.1 |
| ☑ | Parabacteroides distasonis strain BFG-592 chromosome, complete genome | Parabacteroides distasonis | 24255 | 28029 | 36% | 0.0 | 98.54% | 5163710 | CP103256.1 |
| ☑ | Parabacteroides distasonis strain BFG-574 chromosome, complete genome | Parabacteroides distasonis | 24238 | 24331 | 36% | 0.0 | 98.55% | 5298107 | CP103286.1 |
| ☑ | Alistipes finegoldii CE91-St15 DNA, complete genome | Alistipes finegoldii | 23955 | 24058 | 36% | 0.0 | 98.18% | 4117255 | AP025581.1 |
| ☑ | Butyricimonas paravirosa strain DSM 105722 chromosome | Butyricimonas paravirosa | 23941 | 32535 | 36% | 0.0 | 98.25% | 5570268 | CP043839.1 |
| ☑ | Bacteroides eggerthii strain CL11T00C20 chromosome, complete genome | Bacteroides eggerthii | 23859 | 28418 | 36% | 0.0 | 98.04% | 5008416 | CP072227.1 |
| ☑ | Bacteroides uniformis CL03T12C37 chromosome, complete genome | Bacteroides uniformis CL0... | 23752 | 37072 | 57% | 0.0 | 98.01% | 4874733 | CP072255.1 |
| ☑ | Parabacteroides distasonis strain BFG-149 chromosome, complete genome | Parabacteroides distasonis | 23745 | 24187 | 36% | 0.0 | 98.00% | 5335774 | CP103148.1 |
| ☑ | Bacteroides uniformis strain CL06T03C05 chromosome, complete genome | Bacteroides uniformis | 23741 | 28451 | 36% | 0.0 | 97.99% | 5364439 | CP072239.1 |
| ☑ | Parabacteroides distasonis strain BFG-157 chromosome, complete genome | Parabacteroides distasonis | 23739 | 24181 | 36% | 0.0 | 97.99% | 5335766 | CP103222.1 |
| ☑ | Odoribacteraceae bacterium CE91-St21 DNA, complete genome | Odoribacteraceae bacterium | 23736 | 47573 | 36% | 0.0 | 97.98% | 5743229 | AP025578.1 |
| ☑ | Bacteroides thetaiotaomicron strain DFI.6.40 chromosome, complete genome | Bacteroides thetaiotaomicr... | 17101 | 25068 | 36% | 0.0 | 99.43% | 6687652 | CP143941.1 |
| ☑ | MAG TPA_asm: Caudoviricetes sp. isolate ct6ER4, partial genome | Caudoviricetes sp. | 15664 | 26727 | 45% | 0.0 | 95.27% | 23863 | BK038431.1 |
| ☑ | Butyricimonas faecalis strain H184 chromosome, complete genome | Butyricimonas faecalis | 14619 | 37132 | 36% | 0.0 | 97.36% | 4925405 | CP032819.1 |
| ☑ | MAG TPA_asm: Caudoviricetes sp. isolate ctUMx1, partial genome | Caudoviricetes sp. | 10682 | 14522 | 24% | 0.0 | 96.03% | 23200 | BK017507.1 |
| ☑ | Bacteroides uniformis strain BFG-227 chromosome, complete genome | Bacteroides uniformis | 10671 | 27607 | 61% | 0.0 | 96.00% | 4744482 | CP103250.1 |
| ☑ | MAG TPA_asm: Caudoviricetes sp. isolate ctZtD3, partial genome | Caudoviricetes sp. | 10222 | 19872 | 37% | 0.0 | 93.06% | 63305 | BK042668.1 |
| ☑ | MAG TPA_asm: Caudoviricetes sp. isolate ctZfq11, partial genome | Caudoviricetes sp. | 10028 | 33476 | 51% | 0.0 | 95.08% | 68100 | BK047737.1 |
| ☑ | MAG TPA_asm: Caudoviricetes sp. isolate ctVyZ1, partial genome | Caudoviricetes sp. | 9882 | 17672 | 31% | 0.0 | 93.90% | 37458 | BK020418.1 |
| ☑ | MAG TPA_asm: Caudoviricetes sp. isolate ctMrZ4, partial genome | Caudoviricetes sp. | 9864 | 25834 | 45% | 0.0 | 92.13% | 29531 | BK019414.1 |

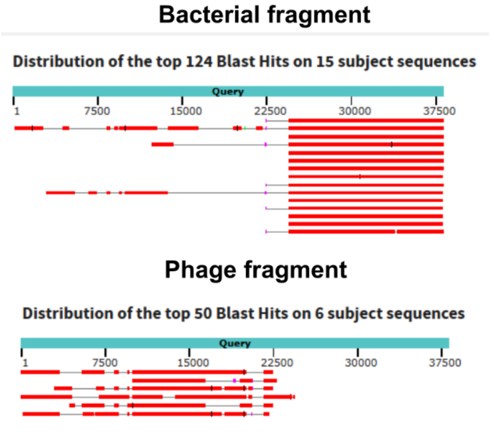

### UHGV-2071483

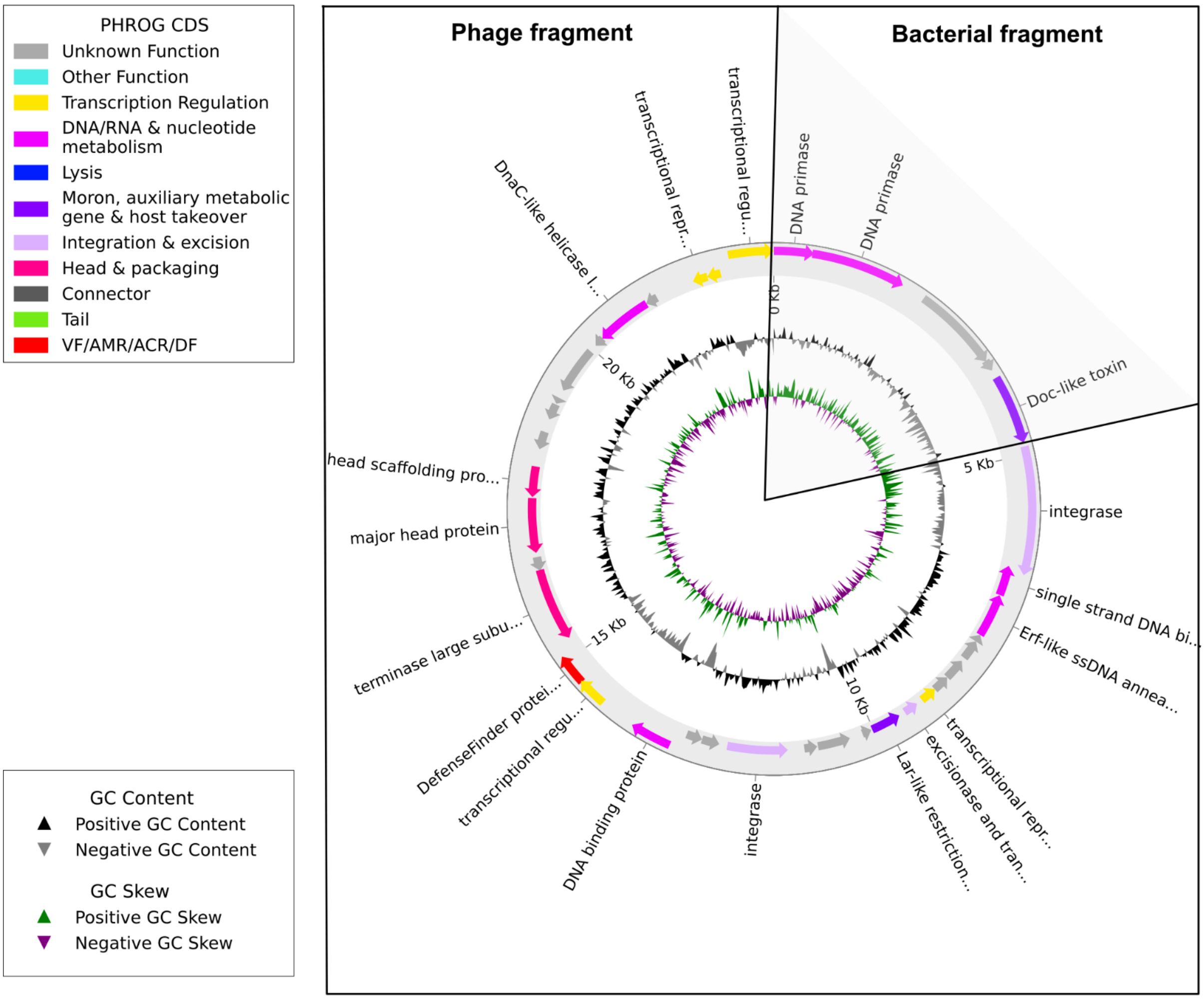


| | Description | Scientific Name | Max Score | Total Score | Query Cover | E value | Per. Ident | Acc. Len | Accession |
|---|---|---|---|---|---|---|---|---|---|
| ☑ | MAG TPA_asm: Caudoviricetes sp. isolate ctk1A4, partial genome | Caudoviricetes sp. | 12443 | 22903 | 64% | 0.0 | 95.49% | 45761 | BK042001.1 |
| ☑ | Faecalibacterium prausnitzii strain UT1 chromosome, complete genome | Faecalibacterium prausnitzii | 12035 | 12319 | 28% | 0.0 | 99.97% | 3135029 | CP170812.1 |
| ☑ | Faecalibacterium prausnitzii SL3/3 draft genome | Faecalibacterium prausnitzii SL3/3 | 12013 | 13447 | 30% | 0.0 | 99.91% | 3214418 | FP929046.1 |
| ☑ | MAG TPA_asm: Caudoviricetes sp. isolate ctfXO8, partial genome | Caudoviricetes sp. | 8757 | 8757 | 21% | 0.0 | 100.00% | 11720 | BK025321.1 |
| ☑ | Hoskinsella hominis strain New-5 chromosome, complete genome | Hoskinsella hominis | 8754 | 8754 | 28% | 0.0 | 90.96% | 3879051 | CP060636.1 |
| ☑ | Faecalibacterium sp. IP-1-18 chromosome, complete genome | Faecalibacterium sp. IP-1-18 | 7023 | 12890 | 30% | 0.0 | 99.97% | 3038545 | CP094472.1 |
| ☑ | Faecalibacterium sp. i25-0019-C1 DNA, complete genome | Faecalibacterium sp. i25-0019-C1 | 7009 | 13424 | 43% | 0.0 | 93.61% | 2876947 | AP031428.1 |
| ☑ | MAG TPA_asm: Caudoviricetes sp. isolate cte129, partial genome | Caudoviricetes sp. | 6872 | 6872 | 20% | 0.0 | 93.46% | 9528 | BK021362.1 |
| ☑ | MAG TPA_asm: Caudoviricetes sp. isolate ctgcl10, partial genome | Caudoviricetes sp. | 6783 | 7324 | 22% | 0.0 | 93.11% | 8197 | BK042947.1 |
| ☑ | MAG TPA_asm: Caudoviricetes sp. isolate ctYbP4, partial genome | Caudoviricetes sp. | 5461 | 5922 | 19% | 0.0 | 91.90% | 12240 | BK056533.1 |
| ☑ | MAG TPA_asm: Caudoviricetes sp. isolate ctiCJ9, partial genome | Caudoviricetes sp. | 5459 | 12040 | 33% | 0.0 | 97.19% | 23686 | BK026048.1 |
| ☑ | MAG TPA_asm: Caudoviricetes sp. isolate ctmwY5, partial genome | Caudoviricetes sp. | 5422 | 6230 | 20% | 0.0 | 91.78% | 42850 | BK026246.1 |
| ☑ | MAG TPA_asm: Caudoviricetes sp. isolate ctZVJ1, partial genome | Caudoviricetes sp. | 5417 | 5772 | 18% | 0.0 | 91.66% | 13057 | BK057962.1 |
| ☑ | MAG TPA_asm: Caudoviricetes sp. isolate ct5Bc6, partial genome | Caudoviricetes sp. | 5363 | 5363 | 17% | 0.0 | 91.46% | 8220 | BK055808.1 |
| ☑ | MAG TPA_asm: Caudoviricetes sp. isolate ctukd10, partial genome | Caudoviricetes sp. | 5228 | 6031 | 18% | 0.0 | 93.64% | 7220 | BK032252.1 |
| ☑ | MAG TPA_asm: Caudoviricetes sp. isolate ctU2p2, partial genome | Caudoviricetes sp. | 5219 | 5219 | 17% | 0.0 | 90.88% | 8846 | BK026374.1 |
| ☑ | MAG TPA_asm: Caudoviricetes sp. isolate ctdLs7, partial genome | Caudoviricetes sp. | 4970 | 7338 | 24% | 0.0 | 89.78% | 9432 | BK028102.1 |
| ☑ | MAG TPA_asm: Caudoviricetes sp. isolate ctHZ63, partial genome | Caudoviricetes sp. | 4887 | 9613 | 28% | 0.0 | 97.26% | 10532 | BK027187.1 |
| ☑ | MAG TPA_asm: Caudoviricetes sp. isolate ctUQP17, partial genome | Caudoviricetes sp. | 4878 | 5934 | 17% | 0.0 | 96.57% | 9760 | BK017540.1 |
| ☑ | MAG TPA_asm: Bacteriophage sp. isolate ctOv11, partial genome | Bacteriophage sp. | 4817 | 5388 | 17% | 0.0 | 91.42% | 9648 | BK025342.1 |

#### Bacterial fragment

**Distribution of the top 14 Blast Hits on 4 subject sequences**

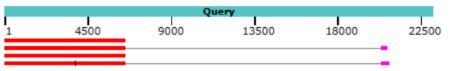


#### Phage fragment

**Distribution of the top 27 Blast Hits on 10 subject sequences**

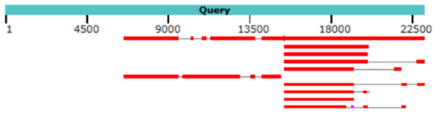

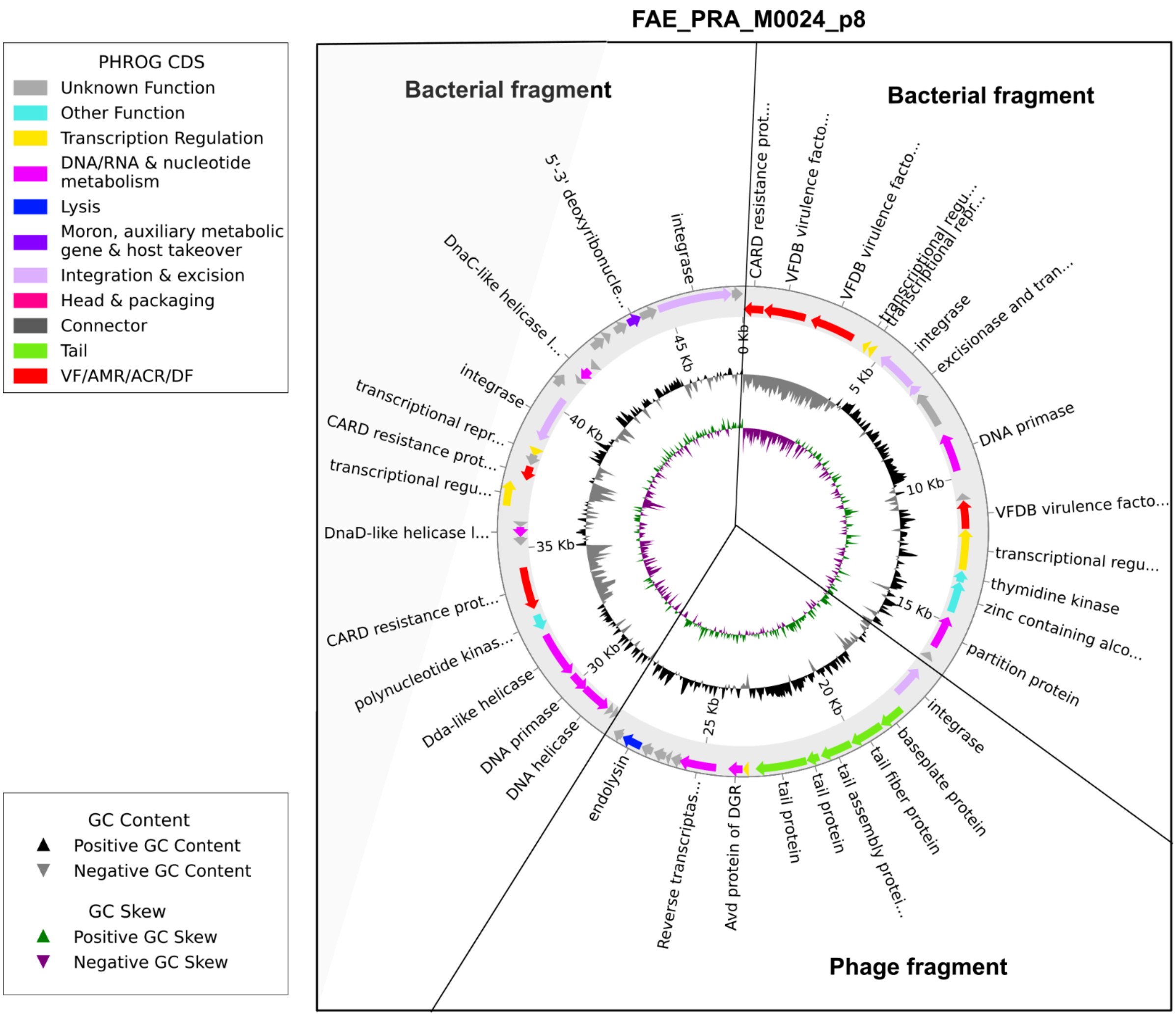


| Description | Scientific Name | Max Score | Total Score | Query Cover | E value | Per. Ident | Acc. Len | Accession |
|---|---|---|---|---|---|---|---|---|
| MAG: Oscillospiraceae bacterium isolate KR001_HIC_0044 chromosome, complete genome | Oscillospiraceae bacterium | 20330 | 67462 | 56% | 0.0 | 97.32% | 2819195 | CP107214.1 |
| Faecalibacterium sp. i25-0019-C1 DNA, complete genome | Faecalibacterium sp. i25-001... | 20131 | 42463 | 42% | 0.0 | 97.02% | 2876947 | AP031428.1 |
| Faecalibacterium prausnitzii strain Fp360 chromosome, complete genome | Faecalibacterium prausnitzii | 19981 | 66051 | 60% | 0.0 | 96.79% | 3039229 | CP065376.1 |
| Faecalibacterium hominis (ex Afrizal et al. 2022) strain 13 chromosome, complete genome | Faecalibacterium hominis (e... | 19130 | 51832 | 50% | 0.0 | 96.85% | 3080408 | CP186860.1 |
| MAG: Bacteriophage sp. isolate 0469_70349, partial genome | Bacteriophage sp. | 18306 | 18639 | 23% | 0.0 | 97.61% | 40135 | OP073258.1 |
| MAG: Bacteriophage sp. isolate 3678_3557, partial genome | Bacteriophage sp. | 18227 | 18552 | 23% | 0.0 | 97.44% | 56148 | OP075713.1 |
| Faecalibacterium prausnitzii strain 28 chromosome, complete genome | Faecalibacterium prausnitzii | 18011 | 49205 | 56% | 0.0 | 99.93% | 2884781 | CP181368.1 |
| Streptococcus suis strain ID48908 chromosome, complete genome | Streptococcus suis | 18005 | 18005 | 21% | 0.0 | 99.93% | 2387659 | CP135090.1 |
| MAG: Bacteriophage sp. isolate 1443_29440, partial genome | Bacteriophage sp. | 18004 | 25098 | 31% | 0.0 | 97.08% | 55841 | OP073925.1 |
| Uncultured bacterium clone 406 genomic sequence | uncultured bacterium | 17983 | 17983 | 21% | 0.0 | 99.94% | 12519 | MW147561.1 |
| MAG: Bacteriophage sp. isolate 1723_80211, partial genome | Bacteriophage sp. | 17581 | 17877 | 23% | 0.0 | 96.37% | 55488 | OP074154.1 |
| MAG: Bacteriophage sp. isolate 2999_112218, partial genome | Bacteriophage sp. | 17285 | 17946 | 22% | 0.0 | 97.82% | 50542 | OP075111.1 |
| MAG: Bacteriophage sp. isolate 2556_50075, partial genome | Bacteriophage sp. | 17173 | 17895 | 23% | 0.0 | 95.73% | 56978 | OP074699.1 |

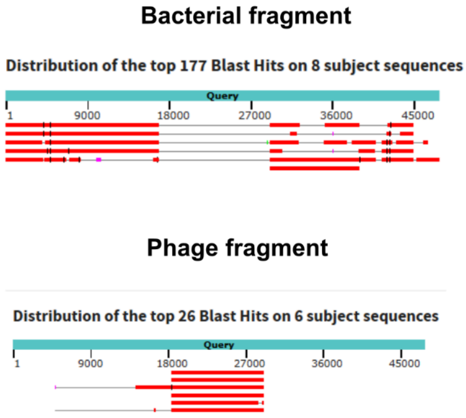


**Figure S12. Bacterial–phage chimeric genome sequences were found to contaminate the reference database**. The most prevalent sequences were annotated with Phold and, together with BLAST results, supported the decision to exclude these sequences from downstream analyses
