## Supplementary Methods for "Cosmopolitan gut bacteriophages expand the phenotype of health-related bacteria"

#### Description

*Faecalibacterium duncaniae* A2-165 was incubated, and extracellular vesicles (EVs) were isolated from the cell-free supernatant for comparative proteomic analysis.

#### Sample Processing Protocol (how to obtain the EVs)

The bacterial strain used in this study was *Faecalibacterium duncaniae* A2–165 (DSM 17677), originally isolated from human faecal material[1]. Cultivation was carried out in a nutrient-rich, complex medium optimised to meet the growth requirements of fastidious *Faecalibacterium* species[2].

Strain *F. duncaniae* A2-165 was grown overnight in liquid medium starting from a single isolated colony on agar plates. The following day, bacterial cells were enumerated by microscopy. An appropriate volume of culture was collected and centrifuged at 3,500 × g for 10 min. The supernatant was discarded, and the bacterial pellet was resuspended in particle-free medium to a final concentration of $8 \times 10^4$ bacteria/mL. The medium was rendered particle-free prior to use by tangential flow filtration (TFF) with a 100 kDa molecular weight cutoff (MWCO) polyethersulfone (PES) membrane (MicroKros®, Repligen).

After incubation, cultures were centrifuged at 3,500 × g for 10 min at 4 °C to remove bacterial cells. The resulting supernatant was filtered through a 0.22 µm pore-size filter on ice. The filtered supernatant was then diluted tenfold in phosphate-buffered saline (PBS), and EVs were isolated and concentrated by tangential flow filtration using a 30 kDa MWCO PES membrane (MicroKros®, Repligen). Purified EVs were stored at −80 °C until further use.
Extracellular vesicles were quantified by nano–flow cytometry (NanoFCM).

### Proteomics

#### Tryptic Digestion in gel

Short SDS-PAGE runs (NuPAGE 4–12% Bis–Tris Gel) were performed, allowing proteins (10 µg) to migrate 0.5 cm. Each lane was excised and sent to PAPPSO plateform facilities (http://pappso.inrae.fr/) for proteomic analysis. Briefly, the protein disulfide bridges were reduced by DTT 10 mM for one hour at 56°C and resulting reduced cysteine were alkylated by iodoacetamide (50 mM) for 1h at room temperature into darkness before the step of enzymatic digestion performed by adding 200 ng of sequencing grade modified trypsin (Promega) diluted in 50 mM NH4HCO3 for 18 h at 37°C for each protein sample. Tryptic peptides were recovered by washing the gel pieces twice in 0.5 % TFA-50 % acetonitrile and once in 100 % acetonitrile and the supernatant was evaporated to dryness. The peptides were then resuspended in 200 µL of loading buffer (0.1 % formic acid and 2 % acetonitrile (ACN) in H2O), prior to LC-MS/MS analysis.

#### Peptide LC-MS/MS analysis

LC-MS/MS analysis was performed using a NanoElute2 (Bruker) LC system connected to a timsTOF Pro mass spectrometer (Bruker) equipped with a CaptiveSpray source. Tryptic peptide mixtures (1 µL) were loaded at 50 bar onto a C18 precolumn (Acclaim PepMap, 5 µm, 20 mm x 100 µm i.d.; Thermo Scientific), and separated on a 25 cm x 75 µm analytical column packed with 1.7 µm C18 beads and integrated emitter tip (IonOpticks, Australia), maintained at 50°C, using a constant flow rate of 250 nL min-1. A multi gradient steps begins at 2 min from 2% to 30% buffer B (0.1 % formic acid and 100 % acetonitrile) over 70 min before ramping to 85% buffer B and sustained for 12 min. Mobile phase is then ramped back to 98% buffer A (0.1 % formic acid and 98% water) and sustained for 2 min. The timsTOF Pro was operated in dia-PASEF, using 24 isolation windows allocated to 12 dia-PASEF scans. The MS settings as follows : Mass range 100 to 1700 m/z, 1/K0 = 0.60 to 1.60 V·s/cm2, Ramp time 100.0 ms, Lock Duty Cycle to 100%, Capillary Voltage 1600V, Dry Gas 3 l/min, Dry Temp 180°C. The dia-PASEF MS/MS settings were as follows: Mass range 352.7 to 953.7 Da, 1/K0 = 0.70 to 1.10 (estimated cycle time: 1.38s) and CID collision energy ramping from 20 eV at 1/K0 = 0.70 to 59 eV at 1/K0 = 1.60. The system was calibrated using three ions from Agilent ESI LC/MS tuning mix (m/z, 1/K0 : 622.0289, 0.9848 Vs/cm2 ; 922.0097, 1.1895 Vs/cm2 ; and 1221.9906, 1.3820 Vs/cm2 ).

#### Peptides and proteins identification

Protein identification was performed with DIA-NN (version 1.8) in library-free mode with deep learning-based spectrum prediction. The analysis included in silico digestion with cleavage rules set at K* and R*, allowing for one missed cleavage, a minimum peptide length of 7, and a maximum of 30 amino acids. Fragment ion m/z values were restricted to the range of 200–1800, and precursor ions to 300–1800 m/z with charge states from 1+ to 4+. Carbamidomethylation of cysteines was set as a fixed modification, and N-terminal methionine excision was enabled. FASTA files used for identification included an in-house contaminant protein database and a species-specific *F. duncaniae* reference proteome (NCBI, GCA_002734145). DIA-NN generated an *in silico* spectral library and reanalysed the data, selecting predicted spectra over experimental ones when more reliable. Protein inference was performed using the relaxed grouping strategy. Output included precursor-level quantification matrices filtered at a 1% false discovery rate (FDR). Mass accuracy thresholds were set to 15 ppm for both MS1 and MS2 levels. The report.tsv file were used for relative quantification analysis.

#### Relative quantification of peptides and proteins

First, peptides are assigned to multiple entries in the protein.group column of the report.tsv file was filtered out. After, control quality, normalisation, filtration and statistical analysis were performed using MCQR[3]. Peaks showing instability in retention time (RT standard deviation greater than 30 s) were filtered out. Then, only peaks with RTs between 400 and 4800 seconds were retained. Data were normalised to compensate for global variations between LC-MS runs using the diff.median.RT normalisation method. Only repeatable peptides were included in the analysis. A repeatable peptide was defined as one detected in at least 1 out of 4 samples. Proteins with at least two such peptides were retained for quantification. Missing values were imputed by linear regression according to the values of the other peptides of the same protein. Protein intensities were computed by summing the normalised values of their specific and repeatable peptides.
